# Inferring the Metabolic Objectives of Mammalian Cells via Inverse Modeling of Fluxomics and Metabolomics

**DOI:** 10.1101/2025.09.23.677837

**Authors:** James Morrissey, Mariana Monteiro, Michael Betenbaugh, Cleo Kontoravdi

## Abstract

Metabolism reflects evolutionary priorities that govern how cells allocate resources. In mammalian cells, metabolic objectives are layered and context-dependent, making it difficult to pinpoint the priorities that underlie observed phenotypes. Here, we introduce ObjFind-M, an inverse optimization framework that infers reaction-level metabolic objectives in mammalian cells directly from fluxomic and metabolomic data. Using Chinese hamster ovary (CHO) cells as a data-rich mammalian cell system, ObjFind-M consistently identifies mitochondrial ATP synthase as the central metabolic driver, supported by key TCA cycle and electron transport chain nodes. Priorities adjust with cellular state, favoring glycolysis-TCA coupling in the growth phase and shifting toward oxidative phosphorylation and redox balance when proliferative activity slows. High recombinant protein producing CHO cells emphasize citrate shuttling and beta-oxidation, linking energy supply with biosynthetic capacity for protein secretion. Benchmarking against conventional objectives demonstrates that maximizing ATP production most accurately reproduces experimental fluxes. By quantifying metabolic objectives directly from data rather than assuming them *a priori*, ObjFind-M provides a framework for identifying reaction-level strategies that shape cellular decision making.

**Graphical Abstract:** 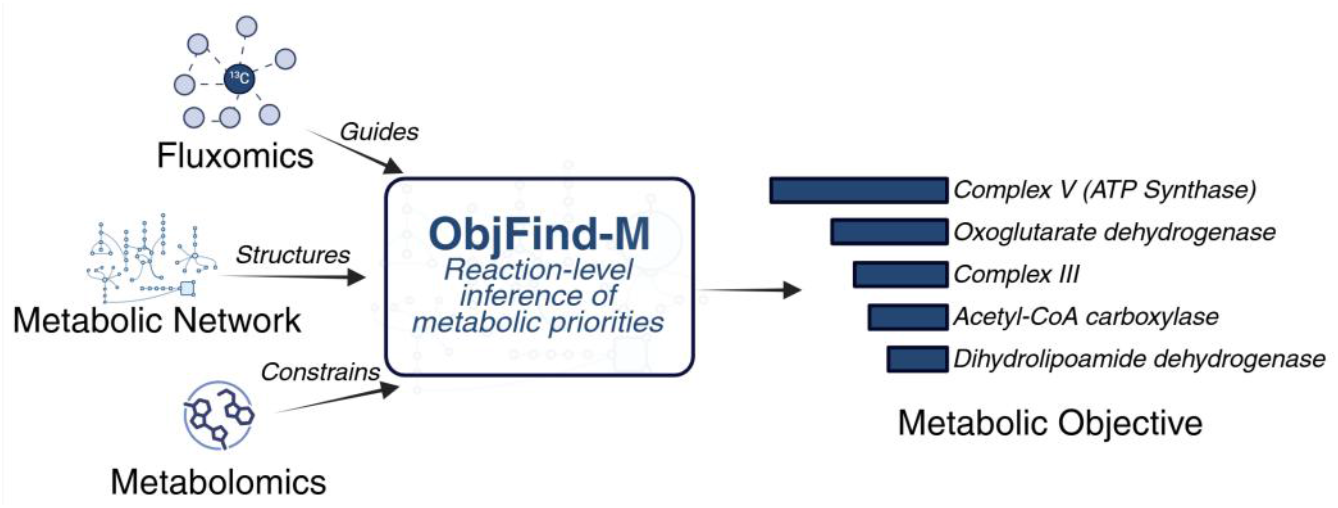

**Highlights:**

- ObjFind-M, an inverse FBA framework, infers reaction-level metabolic objectives from fluxomic and metabolomic data.
- ATP synthase (Complex V) emerges as the top reaction, supported by NADH-generating TCA reactions such as the α-ketoglutarate dehydrogenase complex.
- CHO cells suppress formation of toxic or wasteful byproducts while emphasizing energy-efficient flux routing.
- Growth vs. non-growth phases reveal shifts between biosynthesis and oxidative phosphorylation.
- High recombinant protein producers emphasize citrate shuttling and beta-oxidation, linking energy with secretory capacity.
- ATP demand maximization provides the most accurate generalizable FBA objective

## Introduction

Metabolism captures the cellular priorities that have been shaped by evolutionary pressures, appearing as distinct metabolic “objectives” ^1,2^. These objectives control how a cell coordinates its internal reactions and adjusts to changing physiological or environmental conditions to support growth, maintenance, and specialized functions. Since cells operate within a space of finite resources and physio-chemical constraints, it is not possible to simultaneously optimize all these objectives, leading to trade-offs emerging between competing demands^3^. Uncovering the dominant objective(s) therefore reveals which pathways a cell favors, how it responds to external cues, and where it allocates its limited resources.

The term “objective” in this study is defined as a mixture of evolutionary pressures and external factors that steer cellular behavior at a given time^4^. In simpler organisms, such as bacteria, the predominant objective is typically to maximize growth rate on the available nutrients, which confers a competitive advantage ^5,6^ In contrast, mammalian cells pursue multifaceted objectives within multicellular systems that are not directly associated with single-cell fitness ^7^. For example, astrocytes act as energy reservoirs for neurons ^8^, fast-twitch muscle fibers prioritize rapid ATP production^9^ and hepatocytes prioritize whole-body metabolic homeostasis^10^.

The mathematical representation of metabolic objectives can describe how cells manage limited resources to achieve these biological objectives within mechanistic and environmental constraints ^2^. Systems biology tools are well placed to understand, quantify and contextualize these metabolic objectives, in particular constraint-based optimization approaches, which have been applied to interrogate metabolic networks and characterize cellular metabolism ^11,12^. Flux balance analysis (FBA), the most widely used constraint-based method, seeks the flux distribution that best satisfies an assumed metabolic objective under a given set of constraints ^13^. This objective is the strongest determinant of model accuracy in mammalian cell culture ^14^. However, generic assumptions such as biomass maximization obscure the layered, context-specific priorities that govern metabolism^15–17^. Grounding objective function choice in an evidence-based manner is therefore essential to capture the true metabolic priorities, enabling deeper insight into how mammalian cells allocate energy and resources in health, disease, and bioproduction.

Constraint-based modeling, particularly with genome-scale models (GEMs), has already proven valuable to describe mammalian cell metabolism^18–21^. However, the lack of understanding of mammalian cell metabolic objectives, undermines the predictions using FBA and GEMs, where the *in silico* intracellular fluxes predicted may not reflect *in vivo* behavior. The traditional approach in constraint-based modeling is to use experimental data to constrain a network, then optimize for an assumed metabolic objective function to obtain predicted fluxes (i.e. FBA). Instead, an “inverse” model would use measured data to infer the underlying metabolic objective that best recovers experimental fluxes. This follows the approach first demonstrated in ObjFind ^22^ which was applied to *E. coli*. Since this original method, other inverse optimization approaches have been developed ^23,24^, as well as metabolic task ranking using omics data ^17,25,26^, as summarized in a recent review ^2^. Despite this progress, no existing method has uncovered metabolic goals in mammalian cells at reaction-level resolution using an inverse optimization approach via fluxomic and metabolomic data. This gap limits both the capabilities of model predictions and our understanding of the fundamental strategies shaping mammalian cell physiology.

In this work we develop ObjFind-M (Figure 1), a bilevel optimization algorithm which discovers the metabolic objectives that best describe underlying experimental data in mammalian cells. In the inner level, metabolomics, in the form of exchange rates, set boundary conditions on nutrient uptake and byproduct secretion in an FBA model. In the outer layer, intracellular fluxomics, growth rate, and recombinant protein productivity guide the inferred objective so that model predictions match observed phenotypes. ObjFind-M returns a cellular fitness function, a weighted combination of biochemical reactions that best reproduce the data. These weights, called coefficients of fitness (*CoF*), provide a ranked list of reactions that quantify each reaction’s contribution to the inferred objective. We demonstrate the approach in Chinese hamster ovary (CHO) cells as a model mammalian system, capitalizing on their rich fluxomic and metabolomic datasets that are suitable for discovery of metabolic objectives at the level of individual biochemical reactions.

**Figure 1:**
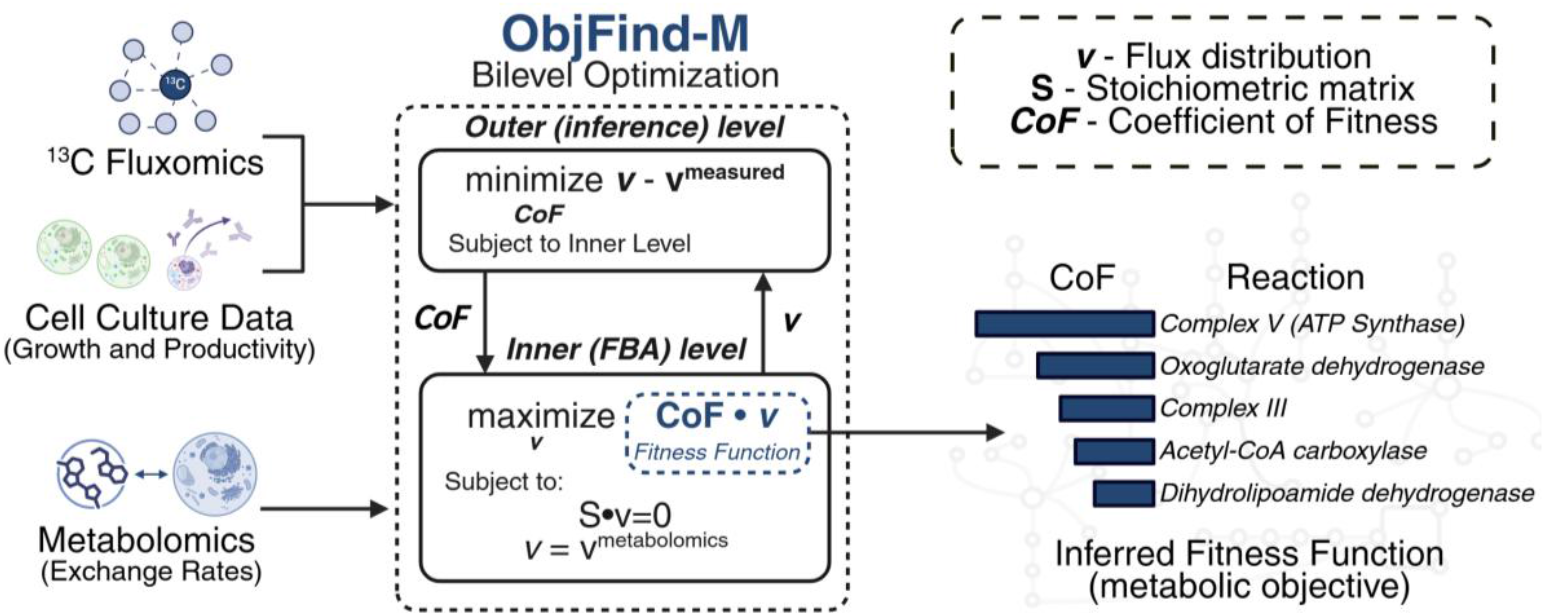
Overview of ObjFind-M methodology. Metabolomics, fluxomics, cell growth and recombinant protein productivity are used as inputs to a bilevel optimization problem. The bilevel optimization finds the linear combination of reactions (fitness function), which, when maximized in an FBA problem, fits the observed experimental data. The output fitness function consists of coefficients of fitness (CoF) for each reaction, revealing which reactions are prioritized in the given condition. Created with BioRender.com.

We first demonstrate the approach on a small-scale, interpretable CHOmpact model ^27^ and then scale to the iCHO2441 GEM ^28^, analyzing a variety of culture conditions. Across datasets and models, the inferred objectives align on a desire to maximize ATP synthase (Complex V), demonstrating that mammalian cells prioritize ATP yield as a central strategy to meet energetic and biosynthetic demands. We then quantify how inferred fitness functions shift between cell growth phases, and identify features associated with high-recombinant protein production. Finally, we benchmark the discovered objectives against commonly assumed objectives in mammalian cells and show that maximizing total ATP demand provides a simple, general objective that aligns FBA predictions with cell behavior. Together, ObjFind-M reveals the metabolic priorities of CHO cells at the biochemical reaction-resolution, providing a more realistic account of cellular physiology and a framework to identify the driving forces that guide cellular metabolism. We present the methodology and code for ObjFind-M to be implemented in other cell systems.

## Results and discussion

### ATP synthase emerges as the key objective node in core metabolic model

To investigate the evolved metabolic priorities of mammalian cells, we developed ObjFind-M, a bilevel inverse optimization framework that infers metabolic objectives directly from fluxomic and metabolomic data (Figure 1). Unlike conventional FBA, which optimizes an assumed objective such as biomass, ObjFind-M identifies the weighted combination of reaction fluxes (termed the *fitness function*) that best reconciles model predictions with observed phenotypes. The resulting coefficients of fitness (*CoF*) quantify how strongly each reaction contributes to the inferred cellular objective, allowing us to reveal context-dependent metabolic priorities without imposing assumptions *a priori*.

We first applied ObjFind-M to CHOmpact ^27^, a small-scale and interpretable model of CHO cell metabolism containing 144 reactions and 111 metabolites. Figure 2 displays the reactions with the highest CoF, averaged across all experiments. The highest contributing reaction is **F19** (NADH → 2.5 ATP + NAD^+^), representing oxidative phosphorylation (OXPHOS) through Complex I and V (ATP Synthase) of the electron transport chain (ETC). This step links NADH generated by glycolysis and the TCA cycle to proton-driven ATP synthesis, making it the core node of energy production. A similarly high coefficient is observed for **F18**, which captures ATP generation via FADH_2_ oxidation through Complex II and ATP synthase. The prominence of both F18 and F19 indicates a cellular strategy to maximize electron flow through the ETC to achieve high ATP yield, with ATP synthase emerging as the central reaction in this objective. This focus on efficient energy conversion reflects evolutionary pressure to extract maximum energy from limited nutrients, ensuring energy supply under diverse conditions ^29^, and in CHO cells, this energy-centric objective aligns with the high biosynthetic and secretory demands biopharmaceutical production ^30,31^. ObjFind-M also assigns weight to **F104**, reflecting a drive to maximize overall ATP production capacity beyond specific pathway nodes.

**Figure 2:**
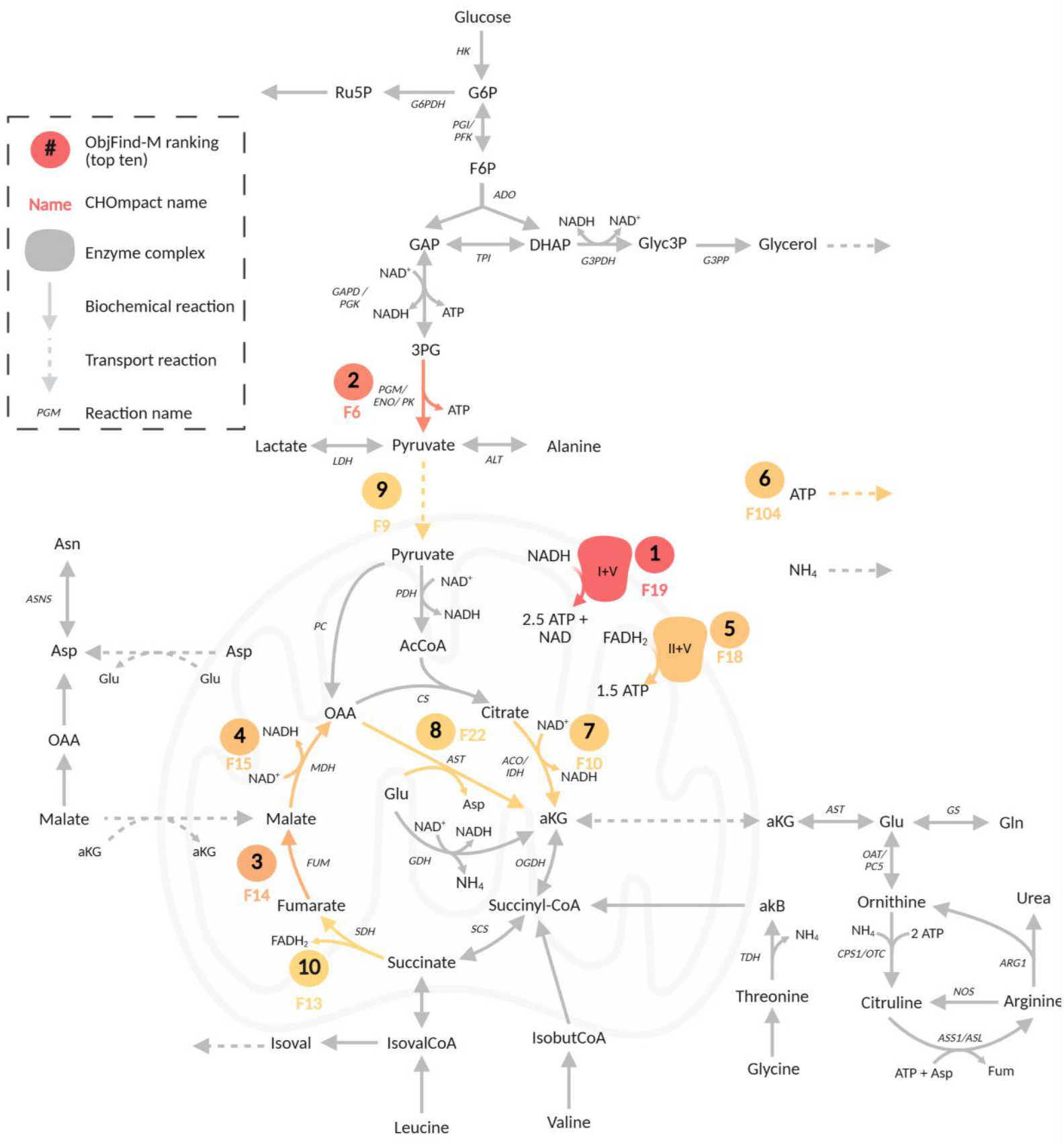
Top ten reactions prioritized in the ObjFind-M-inferred fitness function using the CHOmpact model. Reactions with the highest CoF emphasize oxidative phosphorylation through ATP synthase, glycolysis-derived ATP, and strong TCA cycle activity, highlighting a strategy centered on efficient ATP generation. Created with BioRender.com.

The second most prioritized reaction is **F6**, a lumped phosphoglycerate mutase (PGM), enolase (ENO), and phosphoglycerate kinase (PK) reaction, the final step of glycolysis. The prominence of this step indicates that cells maintain a strong cytosolic ATP supply through glycolysis, complementing the high-yield ATP generated via OXPHOS. By routing carbon through this regulated glycolytic step ^32^ and maximizing pyruvate transport into mitochondria (**F7**), CHO cells sustain a tightly coupled glycolysis–TCA–ETC axis, with nearly half of total ATP reportedly derived from glucose ^33^. At first glance, this reliance on efficient respiration may seem at odds with the Warburg effect, often considered a hallmark of mammalian cells in culture. However, lactate production could be understood as a consequence of proteome allocation limits ^34,35^ or redox balancing requirements ^36,37^ rather than as an intrinsic metabolic objective. Under the tested conditions, mitochondrial respiration consistently emerges as the dominant strategy.

High coefficients for TCA cycle reactions catalyzed by fumarase (**F14**), malate dehydrogenase (**F15**), isocitrate dehydrogenase (**F10**) and succinate dehydrogenase (**F13**) highlight the importance of maintaining high flux through the cycle. These reactions support continuous NADH and FADH_2_ production for OXPHOS, ensuring electron delivery to ETC. In addition, the malate aspartate shuttle (MAS) reaction **F22**, the mitochondrial aspartate transaminase that supports the MAS, receives a substantial coefficient. The MAS transports reducing equivalents from NADH produced in the cytosol into the mitochondria for use in ETC ^38^.

### Core metabolic model reveals minimization of toxic and wasteful byproducts

ObjFind-M not only identifies reactions that dominate cellular objectives, but also the pathways that cells deprioritize to fit experimental behavior. A consistent suppression of pathways that generate toxic or energetically wasteful byproducts (Figure 3) is highlighted, with ammonia export (**F108**) as the most deprioritized reaction. Ammonia is a well-known toxic and inhibitory byproduct in cell culture ^39^, and its minimization suggests a cellular strategy to avoid stress induced by this byproduct. Instead, cells prioritize the urea cycle reactions (**F57, F58, F59, F141**) within the top 20 reactions, as seen in Supp. Data S3. These reactions are maintained even at an ATP cost, highlighting that cells invest energy in detoxifying nitrogen rather than releasing ammonia directly. Cells also appear to deprioritize glutamate dehydrogenase (**F23)** potentially due to this reaction being a large source of ammonia in cell culture ^40^.

**Figure 3:**
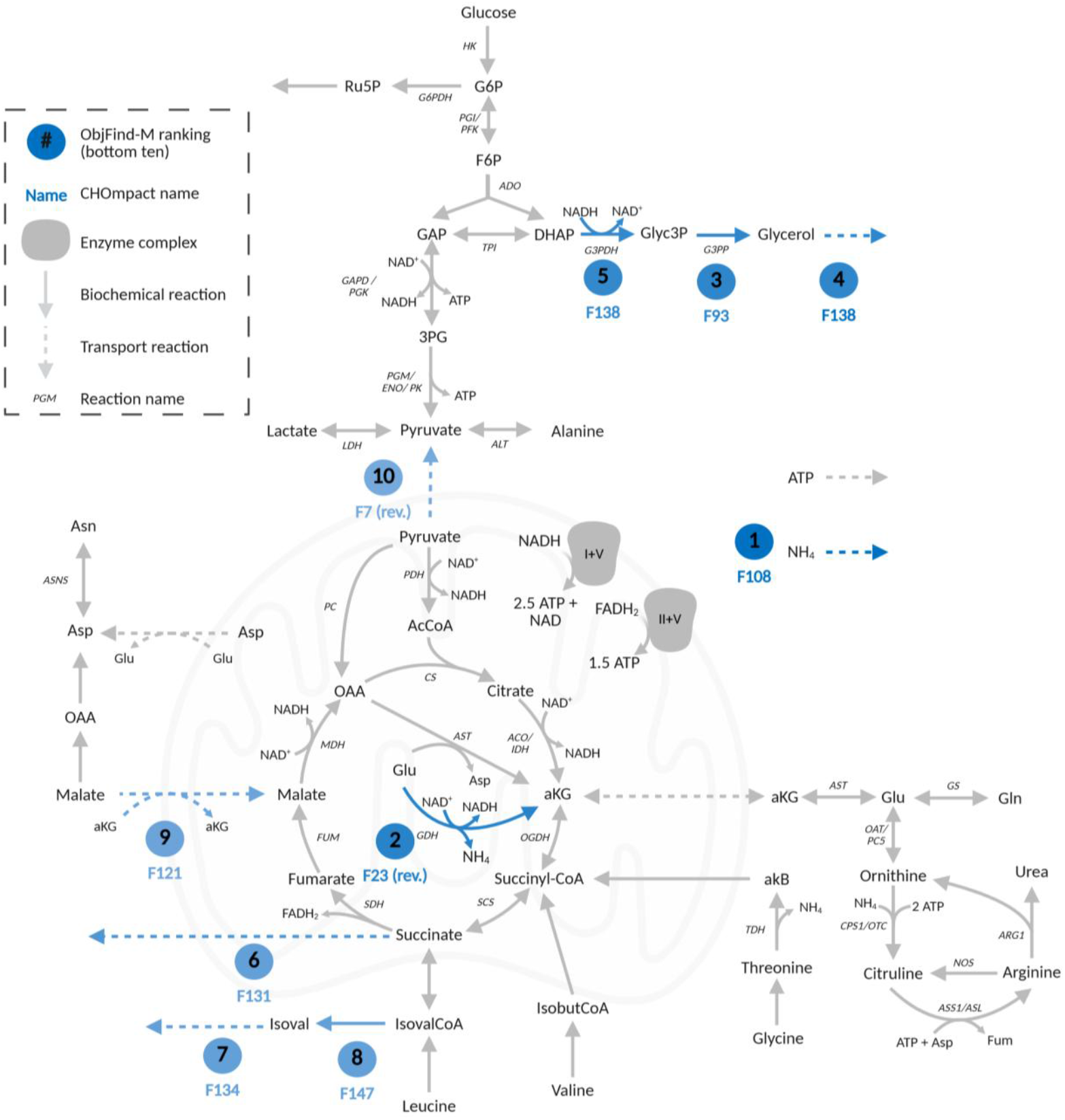
Reactions with the lowest CoF in the ObjFind-M fitness function from the CHOmpact model. CHO cells minimize flux through pathways leading to toxic or wasteful byproducts, including ammonia, glycerol, branched-chain amino acid catabolism, and succinate secretion. This reflects a strategy of reducing metabolic inefficiencies and stress. Created with BioRender.com.

Three reactions with low *CoF* are involved in glycerol formation and export (**F92, F93, F138**). These reactions remove triose phosphates from glycolysis while consuming cytosolic NADH and yielding no ATP. Their uniformly low coefficients indicate that cells treat glycerol formation as a wasteful sink for both carbon and reducing power ^41,42^, potentially preferring to route key intermediates such as glycerol-3-phosphate for lipid biosynthesis. The glycerol phosphate shuttle transfers cytosolic reducing equivalents into the mitochondria by oxidizing cytosolic NADH and reducing mitochondrial FAD to FADH_2_, feeding electrons directly into the ETC ^43^. However, this pathway is not present in CHOmpact, and therefore this pathway is a wasteful carbon and NADH sink that the model attempts to minimize to fit an overall objective to increase ATP production.

Cells also minimize the catabolism of branched-chain amino acids (BCAA) in reactions **F47** and **F134**. The catabolic breakdown of BCAAs generates potentially harmful intermediates, including short-chain fatty acids and acyl-CoAs ^44^. The low weights assigned to these reactions suggest that mammalian cells may have evolved to limit BCAA degradation as a protective strategy, reducing the risk of metabolic toxicity and reduce stress under bioreactor conditions. Succinate secretion (**F131**) is minimized, indicating tight regulation of the TCA cycle at this branch point ^30^. Similarly, **F7_reverse**, representing the export of mitochondrial pyruvate, is minimized and serves as the mirror image of the highly weighted forward transporter. Retaining pyruvate within the mitochondria maximizes ATP yield and supports efficient coupling of glycolysis with the TCA cycle.

Overall, ObjFind-M applied to CHOmpact shows that cells are geared toward maximizing efficient ATP generation while avoiding toxic or wasteful byproducts. The highest weighted reactions emphasize OXPHOS via ATP synthase, supported by the TCA cycle, ATP/NADH producing glycolytic steps, pyruvate transport and the MAS, all linking upstream metabolism to the ETC for efficient energy production. In contrast, pathways that produce toxic byproducts such as ammonia or divert carbon and reducing power into nonproductive sinks such as glycerol formation, BCAA catabolism and succinate export are minimized. These patterns suggest cells have adapted, both through evolutionary pressures and decades of industrial selection (in the case of CHO cells), to favor metabolic routing that maintains high energy availability, reduces toxic stress and supports the high biosynthetic demands of recombinant protein production.

This analysis is unique because it uses data to uncover the internal metabolic goals that drive cell behavior, rather than using assumed objectives, such as growth rate or productivity, to simulate behavior. While the small-scale CHOmpact model offers insight into core metabolic priorities, it represents only a subset of cellular metabolism. To obtain a more complete view of metabolic objectives, we next applied the ObjFind-M framework to the iCHO2441 GEM for a system-level analysis.

### Genome-scale analysis reinforces ATP synthase as the core driver

To evaluate cultured mammalian cell metabolic objectives at a systems-wide level, we applied the ObjFind-M framework to the iCHO2441 GEM ^28^. GEMs capture a broader range of pathways, which enables identification of both primary metabolic goals and supporting secondary functions that collectively shape behavior. ObjFind-M was first applied across all experiments (Figure 4) to get an overview of cell priorities, with the full results table available in the Supp. Data S4.

**Figure 4:**
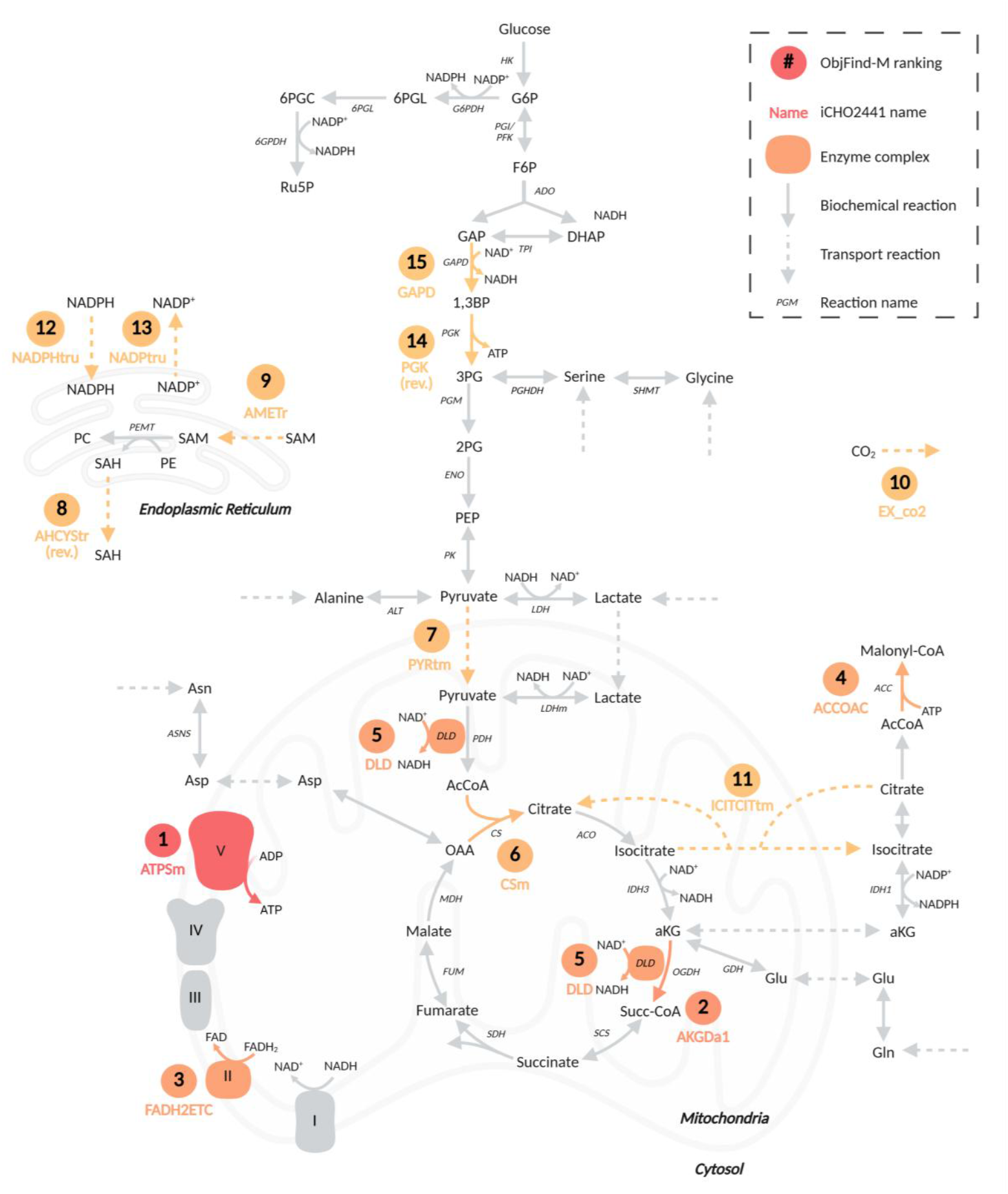
Top 15 reactions prioritized across all experiments in the iCHO2441 genome-scale model. ATP synthase (Complex V) emerges as the dominant objective, supported by TCA cycle reactions, electron transport, and citrate partitioning between mitochondria and cytosol. Additional priorities include lipid biosynthesis and redox balance, reflecting a dual focus on energy and biosynthetic capacity. The fifth highest reaction, **DLD** is a subunit of both the PDH and OGDH complexes and is therefore shown as part of both complexes. Created with BioRender.com.

Figure 4 shows the reactions with the highest CoF in the iCHO2441 GEM. ATP synthase (**ATPS4m)** remains the dominant reaction in the inferred objective, in agreement with the CHOmpact model. In addition, **FADH2ETC**, modeling the transfer of electrons from FADH_2_ to coenzyme Q in Complex II was the third highest scoring reaction, highlighting the underlying goal to maximize ETC flow and mitochondrial ATP generation. Notably, **ATPS4m** appears among the top three reactions in more than half of the 41 datasets analyzed (Supp. Data S4). With more than 8,200 candidate reactions represented in the iCHO2441 network, the repeated prominence of ATP synthase highlights that this node is a central driver of cell metabolism and demonstrates that ObjFind-M can recover physiologically meaningful reaction priorities.

The second most prioritized reaction is **AKGDa1**, representing the E1 component of the oxoglutarate (or α-ketoglutarate) dehydrogenase (OGDH) complex. In cancer biology, OGDH has been recognized as a gatekeeper of OXPHOS^45^, tightly regulating NADH production, TCA flux, and redox balance. That CHO cells elevate OGDH within their inferred objectives may highlight a conserved principle between CHO and cancer cells, where mitochondrial ATP production is safeguarded by maximizing flux through the highly regulated node of the TCA cycle. This finding strengthens the view of respiration as the dominant objective and positions OGDH as a strategic point linking glutaminolysis, where glutamine serves as a major anaplerotic source, with energy supply. Dihydrolipoamide dehydrogenase (**DLD**), the E3 component of the pyruvate dehydrogenase (PDH) and OGDH complexes ^46^, also ranks highly. Together with **AKGDa1**, this highlights that CHO cells also prioritize the regeneration of cofactors required to keep these multi-enzyme complexes running at high capacity. By sustaining NADH output from both glycolytic and glutamine-derived carbon, **AKGDa1**/**DLD** ensures a continuous electron supply to the ETC, reinforcing mitochondrial ATP production as the unifying principle of the inferred objective.

GEM analysis reveals that mammalian cells do not operate with energy generation alone in mind. Citrate synthase **(CSm)** ranks among the highest-contributing reactions, catalyzing the first committed step of the TCA cycle and reinforcing the drive to maintain high mitochondrial flux. Yet citrate’s role extends beyond oxidation in the TCA cycle. The strong weighting of **ICITCITtm**, which mediates citrate/isocitrate transport between mitochondria and cytosol, indicates that cells strategically regulate citrate localization to balance energy production with biosynthesis. A similar principle is observed in cancer metabolism, where glutamine-derived α-KG replenishes isocitrate and citrate to support lipid synthesis under hypoxia or ETC impairment^45^. Once exported, citrate is cleaved by ATP citrate lyase to form acetyl-CoA, a precursor for fatty acid and lipid synthesis ^47^. This is further supported by the prioritization of acetyl-CoA carboxylase **(ACCOAC)**, which converts acetyl-CoA to malonyl-CoA to initiate fatty acid synthesis. Together, these findings reflect an anabolic strategy in which citrate shuttling ties mitochondrial energy metabolism to lipid biosynthesis, supporting membrane expansion and the secretory apparatus.

Supporting ATP synthesis from the glycolytic arm, phosphoglycerate kinase (**PGK_reverse**) and glyceraldehyde-3-phosphate dehydrogenase (**GAPD**) appear among the top reactions, showing that while cells favor mitochondrial respiration, glycolysis remains an important feeder pathway for energy and precursor generation.

S-adenosylmethionine (SAM) transport into the endoplasmic reticulum (ER) (**AMETr**) and S-adenosylhomocysteine (SAH) transport out of the ER (**AHCYStr_reverse**) are both assigned high CoFs. In the ER, SAM is involved in phospholipid synthesis, donating its methyl group to form phosphatidylcholine (PC), a major component of membranes ^48^, and becoming SAH in the process. The maximization of SAM import *into* the ER alongside the maximization of SAH *out* of the ER shows that cells prioritize ER membrane synthesis via recycling SAM and SAH between the compartments. To support this, CHO cells fed with SAM exhibit increased cell-specific productivity ^49^, supporting the idea that enhanced PC synthesis directly contribute to the ER’s ability to accommodate high secretory capacity.

Overall, the inferred objective functions portray cells as oriented toward mitochondrial ATP production, while simultaneously ensuring sufficient biosynthetic capacity for lipid generation and protein processing in the ER. The prioritization of ATP synthase, supported by the OGDH complex to sustain continuous NADH supply from both glucose and glutamine, linked with key glycolytic reactions as well as active citrate localization, indicates that cultured mammalian cells make strategic decisions to balance ATP supply and the provisioning of building blocks for biosynthesis. Cell objectives appear tuned to support metabolic efficiency and productive output under nutrient-controlled, high-demand protein biosynthetic environments, rather than simply maximizing biomass.

### Dual price analysis reveals the structural criticality of the electron transport chain

To complement the CoF analysis, we also examined the dual price (also known as shadow price) associated with each biochemical reaction’s objective constraint. While CoF highlights the reactions most directly prioritized in the inferred fitness function, the dual price reveals which reactions are structurally essential for achieving a solution consistent with experimental data. Table 1 shows the reactions with the highest dual prices from the KKT-based ObjFind-M formulation, averaged across all experiments. These reactions are not always the ones with the largest CoF, but perturbations in their activity strongly restrict the feasible solution space and disrupt the model’s ability to reproduce observed fluxes.

**Table 1:** Top ten reactions with the highest dual price within ObjFind-M, normalized between 0 and 1, then averaged across each experiment. These structurally critical reactions, including ATP synthase, nucleotide and oxygen transport, and ETC complexes, represent essential constraints that the network cannot bypass while fitting experimental data. For intercompartmental transport reactions, (c) represents cytosol, (m) mitochondria and (r) endoplasmic reticulum.

| Model ID | Formula | Reaction Name | Pathway | Dual price (norm.) |
| --- | --- | --- | --- | --- |
| ATPS4m | $\text{ADP} + 4 \text{H}^+ + \text{Pi} \rightarrow \text{ATP} + \text{H}_2\text{O} + 3 \text{H}^+$ | ATP Synthase (Complex V) | Oxidative Phosphorylation | 0.94 |
| ATPtm | $\text{ADP (c)} + \text{ATP (m)} \rightarrow \text{ADP (m)} + \text{ATP (c)}$ | ATP Transport | Transport | 0.74 |
| CYOR_u10m | $2 \text{Cyt c} + 2 \text{H}^+ + \text{Q10H}_2 \rightarrow 2 \text{Cyt c} + 4 \text{H}^+ + \text{Q10}$ | Complex III | Oxidative Phosphorylation | 0.47 |
| H2Otm_reverse | $\text{H}_2\text{O (m)} \rightarrow \text{H}_2\text{O (c)}$ | Water Transport | Transport | 0.44 |
| EX_o2(e)_reverse | $\rightarrow \text{O}_2$ | Oxygen Export | Transport | 0.33 |
| O2t | $\text{O}_2 \text{ (e)} \rightarrow \text{O}_2 \text{ (c)}$ | Oxygen Transport | Transport | 0.32 |
| Plt2m | $\text{H}^+ \text{ (c)} + \text{Pi (c)} \rightarrow \text{H}^+ \text{ (m)} + \text{Pi (m)}$ | Phosphate/Proton Transport | Transport | 0.31 |
| PPAm | $\text{H}_2\text{O} + \text{PPi} \rightarrow \text{H}^+ + 2 \text{Pi}$ | Pyrophosphate Hydrolysis | Oxidative Phosphorylation | 0.30 |
| Htr | $\text{H}^+ \text{ (c)} \rightarrow \text{H}^+ \text{ (r)}$ | Proton Transport | Transport | 0.27 |
| NADH2_u10m | $5 \text{H}^+ + \text{NADH} + \text{Q10} \rightarrow 4 \text{H}^+ + \text{NAD}^+ + \text{Q10H}_2$ | Complex I | Oxidative Phosphorylation | 0.25 |

The reaction with the highest dual price is ATP synthase (**ATPS4m**), the same reaction with the highest CoF in both iCHO2441 and CHOmpact. This highlights that ATP production via OXPHOS is both favored by the cell’s metabolic landscape and crucial to achieving a flux distribution that fits measured data. ATP synthase is thus both functionally and structurally dominant in mammalian cell metabolism.

Other ETC reactions, including Complex I (**NADH2_u10m**) and Complex III (**CYOR_u10m**), also display high dual prices. These steps are required to maintain electron transfer and proton pumping across the inner mitochondrial membrane, which sustains the proton motive force necessary for ATP synthesis. Their structural importance means that constraining or removing them would cause widespread changes to flux distributions. Oxygen transport reactions also emerge as structurally critical, with both oxygen uptake (**EX_o2(e)_reverse**) and transport to cytosol (**O2t**) carrying large dual prices. Their inclusion highlights the importance of oxygen availability for oxidative metabolism. Without continuous oxygen transfer across compartments, the ETC cannot operate, and the model fails to reproduce measured fluxes.

Reactions related to nucleotide transport and cycling, such as ATP transport between mitochondria and cytosol (**ATPtm**), phosphate/proton transport (**PIt2m**), and proton transport into the ER (**Htr**), also rank highly. These reactions ensure that energy produced in mitochondria is available in the cytosol, that phosphate is accessible for phosphorylation reactions, and that organelle environments are properly maintained for biosynthesis. Their high dual prices indicate that such coupling reactions are essential for maintaining energetic balance and supporting biosynthesis. Several high-ranking reactions contribute to thermodynamic feasibility and removal of reaction byproducts. Water transport from the mitochondria to the cytosol (**H2Otm_reverse**) and mitochondrial pyrophosphate hydrolysis (**PPAm**) help drive reactions forward by removing end products that would otherwise accumulate and impede flux.

### Culture phase dictates metabolic priorities

Next, to understand cellular objectives shift with cellular context, we tested how the phase of cell culture impacts the inferred fitness functions. To do so, we selected six experiments from the same study ^50^, with three datasets in the growth phase and three in the non-growth phase (Figure 5). The full *CoF* dataset can be found in Supp. Data S5.

**Figure 5:**
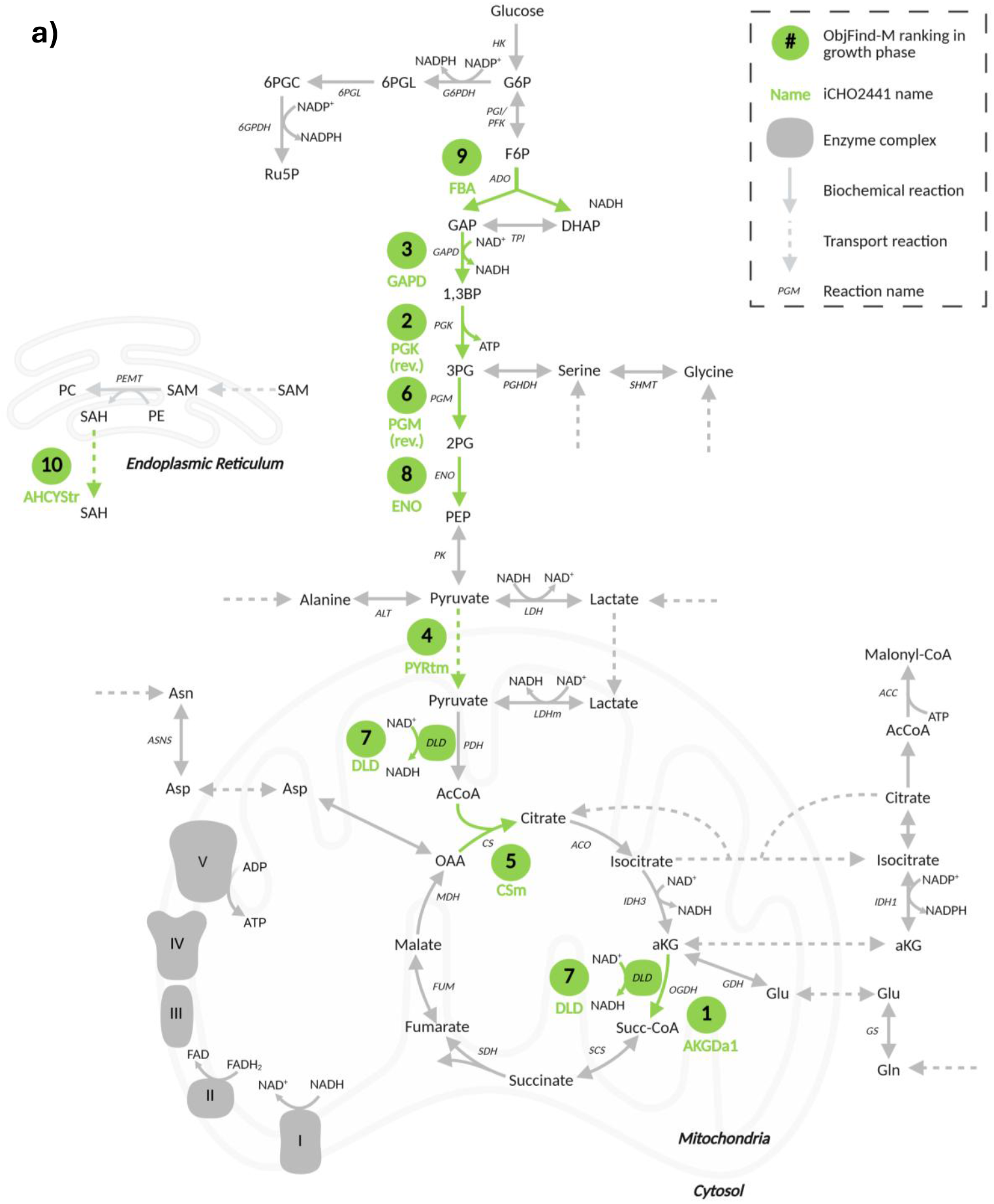
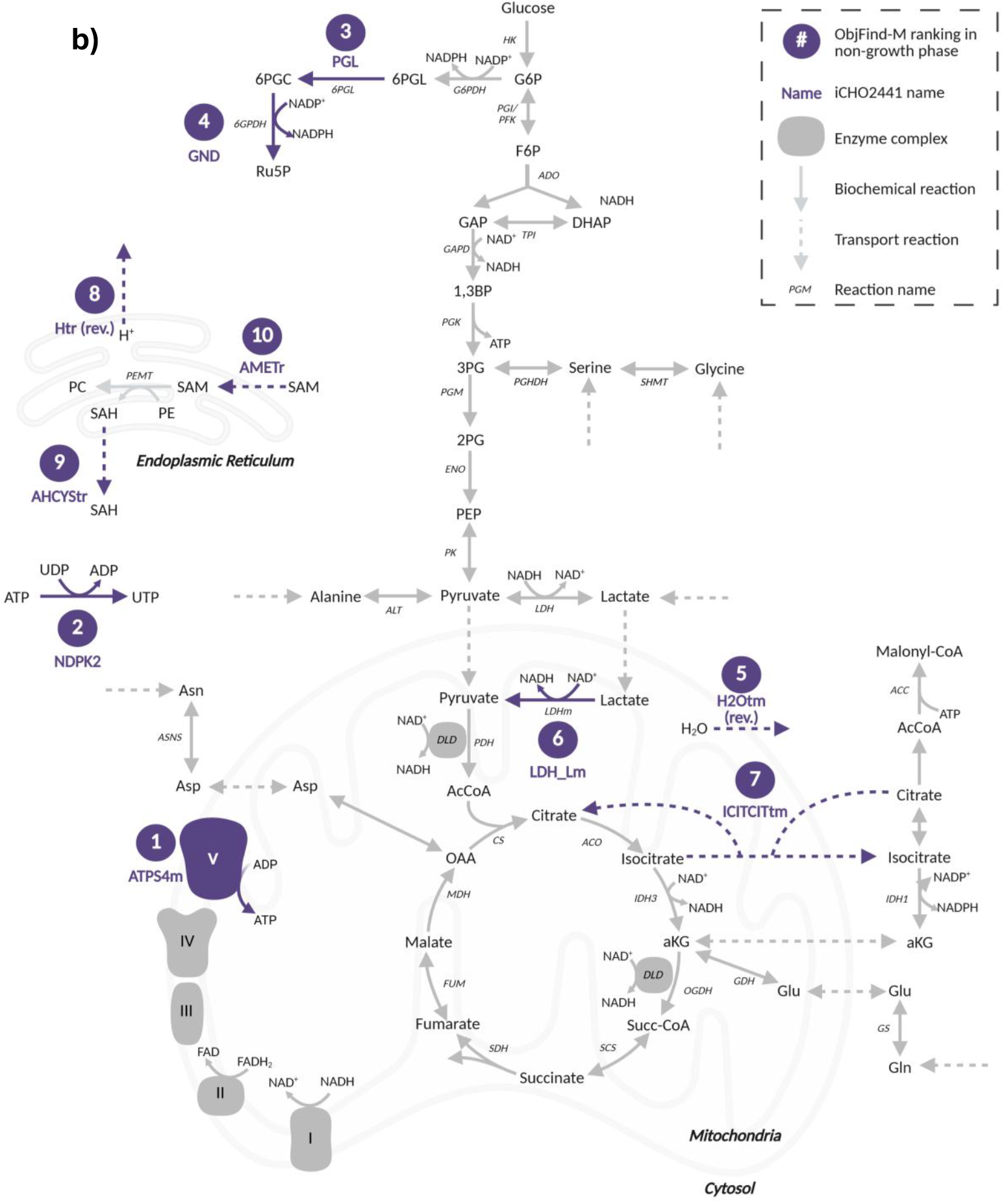
Comparison of inferred metabolic objectives between **a)** growth and **b)** non-growth phase in the iCHO2441 GEM. Top contributing reactions averaged for three growth phase experiments, and three non-growth phase experiments are shown. Exponential-phase cells prioritize glycolysis–TCA coupling to support growth and biosynthesis, while non-growth phase cells simplify their objective toward efficient ATP generation through oxidative phosphorylation, with secondary emphasis on redox balance and nucleotide recycling. Created with BioRender.com.

During the growth phase, the objective is dominated by reactions linking glycolysis and the TCA cycle (Figure 5a). The top-weighted reaction is **AKGDa1**, the E1 component of the OGDH complex, as discussed earlier, which channels glucose and glutamine derived carbon into NADH production for OXPHOS. This priority is reinforced by **DLD**, the E3 subunit of OGDH and PDH complexes, which re-oxidizes the lipoamide cofactor to sustain flux through the complex. Many of the highest-weighted reactions are glycolytic, consistent with observations that CHO cells prioritize rapid glucose uptake in the growth phase ^30^.

The high contribution of pyruvate transport to mitochondria (**PYRtm**), citrate synthase (**CSm**), and **DLD** underscores the strong coordination between glycolysis and mitochondrial metabolism during the growth phase. **PYRtm** channels glycolytic pyruvate into mitochondria, where PDH converts it to acetyl-CoA that enters the TCA cycle via **CSm. DLD**, shared by the PDH and OGDH complexes, sustains NADH generation to feed the ETC and aligns with the prominence of OGDH-related steps noted above. Together these reactions route glucose- and glutamine-derived carbon through glycolysis and the TCA cycle, supporting ATP production and supplying biosynthetic precursors needed for proliferation. This suggests that during the growth phase, a more balanced objective is selected, which considers biosynthetic needs and not just pure mitochondrial respiration.

In the non-growth phase, the inferred metabolic objective is overwhelmingly dominated by ATP synthase (**ATPS4m**), shown in Figure 5b, indicating a shift in cellular priorities toward efficient energy production. As cells exit the proliferative state, their objective function narrows to sustaining ATP generation through OXPHOS, it has been estimated that 80% of ATP produced during this phase is for non-growth-related processes ^33^. Alongside **ATPS4m** other reactions in the objective formulation reflecting metabolic maintenance in the non-growth phase. Among them is **NDPK2**, a nucleotide exchange reaction that regenerates UTP from UDP using ATP. Its inclusion suggests a strategy of nucleotide recycling, supporting essential cellular functions without the burden of de novo nucleotide synthesis ^48^.

Two reactions from the oxidative branch of the pentose phosphate pathway **PGL** (6-phosphogluconolactonase) and **GND** (6-phosphogluconate dehydrogenase), are among the highest contributing reactions in the non-growth phase. Their prioritization indicates that NADPH generation via the PPP is an intrinsic objective of mammalian cell culture during this stage. NADPH is critical for sustaining redox balance and powering antioxidant systems such as glutathione (GSH), used to counter elevated oxidative stress during the non-growth phase ^51^. Selvarasu et al. observed progressive depletion of intracellular GSH/GSSG pools and accumulation of oxidized glutathione (GSSG) extracellularly as culture advanced, underscoring the need to maintain high PPP capacity over time. Supporting this view, PPP fluxes have been reported to increase 5-to 6-fold in the non-growth phase compared to growth, corresponding to ∼30% of glucose uptake ^52,53^.

A unique feature of non-growth phase metabolism captured in the inferred fitness function is the lactate-pyruvate-shuttle (LPS). In this pathway, lactate, either sourced from cytosolic pyruvate via LDH, or taken up from the extracellular environment, is transported into the mitochondria and converted back to pyruvate by mitochondrial LDH (**LDH_Lm**). This process generates NADH directly in the mitochondria to fuel the ETC while also bypassing the bottleneck of pyruvate transport through the mitochondrial pyruvate carrier ^54^. This lactate shuttling phenomenon has been of interest to the metabolic community, since it challenges the notion of lactate simply being a waste product ^55^. This phenomenon has been observed in FBA models ^35,56^ but not yet verified experimentally in cell culture.

### High protein producers prioritize lipid and energy metabolism

To test whether metabolic objectives explain differences in recombinant protein biosynthesis, we compared ObjFind-M–inferred fitness functions between high- and low-producing CHO cells. We analyzed 20 fed-batch experiments producing a monoclonal antibody (Supp. Data S2). Principal component analysis (PCA) of the fitness function coefficients revealed a clear separation between high and low producers along the first principal component (PC1) (Supp. Data S6). This separation suggests that differences in objective function structure are linked to recombinant protein output.

The comparison highlights the strategies that allow cells to sustain high biosynthetic activity, providing insight into how energy and resources are allocated under heavy secretory demand and providing actionable insights for upstream bioprocessing. The reactions identified here are not simply those with the highest fluxes. Instead, they are the reactions that ObjFind-M assigns greater weight in the inferred fitness function of high producers that explain the separation on PC1, demonstrating which internal strategies are associated with high protein producers.

To assess which reactions in the objective function were driving this separation we evaluated the CoFs with the high contribution to the loadings of PC1 (Table 2). The first observation is that productive cells maximize TCA cycle reactions such as **FUMm, DLD** and **CSm**. These reactions reinforce the notion that cells with higher TCA throughput and OXPHOS are associated with higher recombinant protein production ^57–63^. High mitochondrial throughput provides NADH and FADH_2_ to drive OXPHOS to generate ATP for the energy-intensive demands of protein synthesis. Pyruvate metabolism and the PDH complex has previously been identified as a selection marker for high biologics producers ^64–66^, and the appearance of **DLD** (its E3 subunit) in our analysis suggests that these high producers prioritize this step, increasing pyruvate entry into the TCA cycle and acetyl-CoA availability for fatty acid synthesis

**Table 2:** Reactions contributing most strongly to separation of high- vs. low-producing phenotypes in PC1. High producers emphasize TCA cycle activity, citrate shuttling, and beta-oxidation, linking efficient energy supply with biosynthetic capacity. For intercompartmental transport reactions, (c) represents cytosol, (m) mitochondria, (r) endoplasmic reticulum and (x) peroxisome.

| Reaction ID | Reaction Formula | Reaction Name | Pathway |
| --- | --- | --- | --- |
| FUMm | $\text{Fum} + \text{H}_2\text{O} \rightarrow \text{Mal}$ | Fumarase | TCA Cycle |
| ICITCITtm | $\text{Cit (c)} + \text{ICit (m)} \rightarrow \text{Cit (m)} + \text{ICit (c)}$ | Citrate/Isocitrate Antiporter | TCA Shuttle |
| HCO3E | $\text{CO}_2 + \text{H}_2\text{O} \rightarrow \text{H} + \text{HCO}_3$ | Carbonic Anhydrase | pH Regulation |
| CSm | $\text{AcCoA} + \text{H}_2\text{O} + \text{Oaa} \rightarrow \text{Cit} + \text{CoA} + \text{H}^+$ | Citrate Synthase | TCA Cycle |
| FADFADH2tm | $\text{FAD (c)} + \text{FADH}_2 \text{ (m)} \rightarrow \text{FAD (m)} + \text{FADH}_2 \text{ (c)}$ | FAD/FADH<sub>2</sub> Transport | Electron Shuttle |
| DLD | $\text{Dhlm} + \text{NAD}^+ \rightarrow \text{H}^+ + \text{Lpam} + \text{NADH}$ | Dihydrolipoyl Dehydrogenase | Energy Metabolism |
| CO2t_reverse | $\text{CO}_2 \text{ (c)} \rightarrow \text{CO}_2 \text{ (e)}$ | CO<sub>2</sub> Transport | Transport |
| H2Oter_reverse | $\text{H}_2\text{O (r)} \rightarrow \text{H}_2\text{O (c)}$ | H<sub>2</sub>O Transport | Transport |
| Htr | $\text{H}^+ \text{ (c)} \rightarrow \text{H}^+ \text{ (r)}$ | H<sup>+</sup> Transport | pH Regulation |
| OCCOAtm | $\text{Octanoyl-CoA (c)} \rightarrow \text{Octanoyl-CoA (m)}$ | Octanoyl-CoA Transport | Fatty Acid Transport |
| ACOAD120p | $\text{DDCA-CoA} + \text{O}_2 \rightarrow \text{DD2-CoA} + \text{H}_2\text{O}_2$ | Acyl-CoA Dehydrogenase (C12:0) | Beta-Oxidation |
| ACACT140p | $\text{3-OTD-CoA} + \text{CoA} \rightarrow \text{AcCoA} + \text{DDCA-CoA}$ | 3-Ketoacyl-CoA Thiolase (C14:0) | Beta-Oxidation |
| O2tp | $\text{O}_2 \text{ (c)} \rightarrow \text{O}_2 \text{ (x)}$ | O<sub>2</sub> Transport to Peroxisome | Transport |

**CSm** and the citrate/isocitrate antiporter (**ICITCITtm**) appear among the strongest contributors, highlighting the dual role of citrate as both a TCA cycle intermediate and a precursor for cytosolic acetyl-CoA. The ability to shuttle citrate between mitochondria and cytosol allows cells to balance ATP production with lipid biosynthesis for membrane expansion, which is essential to maintain the secretory machinery under high recombinant protein loads. Citrate has been pinpointed in previous studies ^65,67^, suggesting that utilization of this metabolite, either in the TCA cycle or for lipid synthesis, is a hallmark of productive cells.

Reactions involved in fatty acid metabolism, including octanoyl-CoA transport (**OCCOAtm**) and peroxisomal beta-oxidation steps (**ACOAD120p** and **ACACT140p**), as well as oxygen transport to the peroxisome (**O2tp**) contribute strongly to PC1. This suggests high-producing cells favor beta-oxidation as an auxiliary ATP supply route, taking advantage of its high energetic yield compared to glycolysis. The same logic underlies the metabolism of heart cells, which meet their immense energy demands by deriving 50–70% of ATP from fatty acid beta-oxidation ^68^. CHO cells with high productivity have previously been shown to upregulate beta-oxidation genes ^69^. Medium-chain fatty acids like octanoate can enter mitochondria without the carnitine shuttle, enabling rapid conversion to acetyl-CoA, NADH, and FADH_2_ to feed into the TCA cycle and the ETC. These fatty acids are therefore a potential media supplement, however a recent study found that fatty acid supplementation did not significantly increase recombinant protein synthesis ^70^. This suggests that the metabolic features identified here reflect an intrinsic strategy, with high producers hardwired to exploit beta-oxidation as part of their core metabolic objectives rather than relying on external supplementation.

The flavin shuttle between cytosol and mitochondria (**FADFADH2tm**) is also associated with high producers. This reaction keeps mitochondrial flavoproteins oxidized by importing FAD and exporting excess FADH_2_, preventing local redox bottlenecks when ETC flux is high. Because high producers rely on Complex II and mitochondrial beta-oxidation, both are FAD/FADH_2_-dependent; an increased weight on **FADFADH2tm** is consistent with elevated flavin turnover and more efficient electron delivery to the ubiquinone pool. Consistent with this interpretation, high recombinant protein producers have been reported to maintain larger intracellular pools of electron carriers ^57,71^.

The inverse optimization approach presented by ObjFind-M is novel because it uncovers the underlying metabolic priorities that distinguish high producers, rather than merely describing observed phenotypes. By inferring the cellular ‘decision-making process,’ ObjFind-M reveals the strategies cells use to balance energy and biosynthetic demands, enabling us to identify and select for beneficial traits critical to bioproduction.

### What is the appropriate objective function in mammalian cell culture?

A key outcome of this study is to determine the most appropriate metabolic objective function when performing FBA for mammalian cells. The traditional approach is to maximize growth rate (biomass), and while there are a variety of alternative objectives ^72^, it is unclear which of these best reflects cellular behavior. ObjFind-M works by identifying the linear combination of reactions (fitness function) that represents the metabolic priorities under the conditions tested. However, ObjFind-M requires fluxomic and metabolomic data to “rediscover” the objective *de novo* for each condition, which restricts its immediate utility for routine FBA applications. For most users, a simple, universally applicable objective function would be more practical.

We therefore evaluated candidate objectives derived from ObjFind-M’s results alongside other common FBA objectives, by comparing the root mean squared error (RMSE) between ^13^C fluxomic data and predicted fluxes for intracellular reactions. FBA was performed for a variety of candidate objectives, with ^13^C fluxomic data used to benchmark intracellular predictive performance. Flux sampling, an alternative to FBA in which there is no assumed metabolic objective and fluxes are instead sampled randomly from the feasible solution space, was also benchmarked.

Figure 6 shows the RMSE for each candidate objective function, comparing predicted fluxes against ^13^C fluxomics. Not surprisingly, ObjFind-M performs best, since it infers a unique fitness function for each experiment that minimizes the error relative to these measured fluxes. While ObjFind-M is effective at uncovering cellular priorities at the reaction-level, it is not always practical to re-infer a custom objective for every new condition due to the requirement of ^13^C fluxomics. For routine FBA, a more generalizable objective that reflects the key decision-making patterns uncovered in this study is preferable.

**Figure 6:**
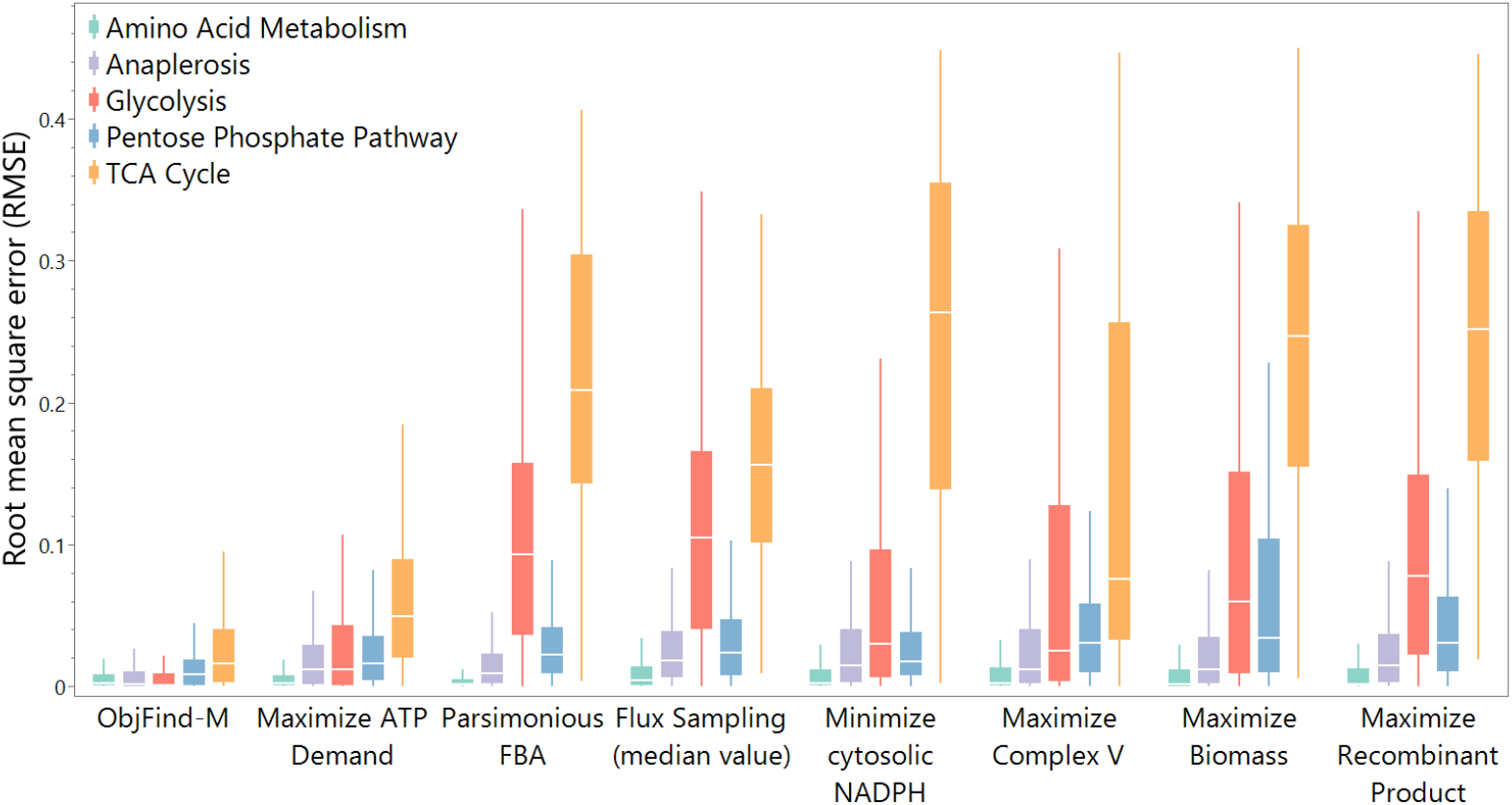
Benchmarking candidate objective functions in iCHO2441 against experimental ^13^C fluxomics. Error metrics (RMSE) quantify how well each assumed objective reproduces observed intracellular fluxes (lower RMSE is better). Maximizing ATP demand outperforms commonly used objectives such as biomass maximization, underscoring ATP-driven energy production as the most realistic general objective for CHO cells. The box represents the interquartile range (IQR), bounded by the first quartile (Q1) and the third quartile (Q3). The whiskers extend to the smallest and largest data points within 1.5 times the IQR from Q1 and Q3, respectively.

The most effective of these generalized objectives is maximizing ATP demand (**DM_atp**), as shown in Figure 6. Optimizing this reaction drives the flux distribution to maximize ATP generation across the entire network. This simple objective substantially reduces error compared to other formulations, particularly improving predictions in glycolysis and the TCA cycle. By balancing fluxes to meet cellular energy needs, it yields intracellular predictions consistent with experimental observations. These results align with ObjFind-M findings, which identified mammalian cells as being metabolically inclined to maximize ATP yield. In the absence of custom objectives that can be fit to the experimental data, it is recommended to select maximization of ATP demand as the metabolic objective of mammalian cells in culture.

Complex V maximization improved TCA and glycolysis predictions when compared to the biomass objective, however not nearly as well as total ATP demand maximization, as demonstrated in Figure 6. This suggests that, while Complex V synthesis is the reaction with the singular highest CoF, it plays a part of a broader cellular strategy to maximize ATP production across all reactions. Maximizing this reaction alone misses the intricacies of ATP production across other pathways and the nuance of other cellular objectives that are better captured by maximizing total ATP demand instead of just ATP synthase alone.

Flux sampling, which does not assume an objective and instead samples the entire solution space, represented a moderate improvement over biomass objective function. However, the key advantage of sampling is to assess a probability distribution for each reaction flux, and taking the median value detracts from this. Flux sampling may be more effective if guided by an appropriate objective to steer the solution space toward biologically relevant states

Other objective formulations did not perform as well. For example, minimizing cytosolic NADPH, an objective previously applied to mammalian cultures ^14^, slightly improved glycolytic flux predictions relative to biomass maximization, but overall was far less effective than maximizing ATP demand. Conventional objectives such as biomass or recombinant protein maximization were the weakest predictors, reflecting the fact that mammalian cells do not operate under a single growth or production driven goal. Instead, their metabolism is shaped by layered and competing priorities, making energy-centered objectives better suited to capture their underlying physiology.

## Conclusion

Metabolic objectives describe how cells allocate resources to achieve their goals within the limits set by biological, physicochemical, and environmental constraints. These objectives, shaped by evolutionary pressures, determine the priorities and trade-offs that govern cellular metabolism ^2^. Systems biology approaches, such as constraint-based modeling, provide a framework to quantify these priorities and simulate cellular behavior, however suffer from limited understanding of metabolic objectives, particularly for mammalian cells which have complex, context-dependent goals. In this study, we develop ObjFind-M, a bilevel inverse-optimization framework to infer objective functions at the biochemical reaction-level directly from fluxomic and metabolomic data.

Using CHO cells as a well-characterized and data-rich mammalian cell system, ObjFind-M infers the prioritization of ATP production through Complex V (ATP synthase). Supporting ATP synthase, key TCA cycle and energy reactions yielding NADH and FADH_2_ are prominent in the metabolic objective, as well as reactions supporting lipid and ER membrane synthesis. Cells appear programmed for metabolic efficiency, rather than simply maximizing biomass. This ATP synthase prominence is uncovered in both a small-scale (CHOmpact) and genome-scale model (iCHO2441). CHOmpact results show low priority is assigned to toxic and wasteful pathways, including BCAA catabolism and the production of ammonia and glycerol, indicating that CHO cells seek to minimize these routes when possible. iCHO2441 analysis reveals the ETC, particularly Complexes I, III and V, is both functionally and structurally critical to mammalian cell metabolism.

Cells reformulate their metabolic goals based on the environmental context and culture conditions. During the growth phase, cells prioritize glycolysis–TCA coupling to feed respiration and supply precursors for growth, then shift toward energy-efficient ATP generation, redox stability, and reliance on lactate shuttling as they enter the non-growth phase. Furthermore, ObjFind-M uncovers different fitness functions between high and low recombinant protein producers, with high producers favoring TCA cycle throughput, beta-oxidation, as well as active localization of citrate to balance between energetic and lipid biosynthesis demands.

Since ObjFind-M reveals that cells adopt an ATP-centric strategy, using total ATP demand as the FBA objective function reproduces experimental fluxes more accurately than classic objectives such as biomass or recombinant protein production. This leads us to propose ATP demand as the preferred objective for constraint-based analyses of CHO cell metabolism.

Applying ObjFind-M to other mammalian systems, including distinct tissues or disease models, may uncover additional context-specific priorities layered on top of the drive for efficient ATP yield. Because the inferred objectives reconcile FBA predictions with experimental observations, they are reliant on the data used to constrain and guide the model. Extending this framework to include additional metabolites or multi-omics layers will likely reveal further nuanced metabolic goals. Nevertheless, our analysis consistently points to ATP synthase and efficient energy metabolism as a core objective.

Overall, ObjFind-M reveals metabolic objectives at the resolution of individual biochemical reactions, enabling quantification of the key forces that shape decision making in cell metabolism. In CHO cells, the method reveals an internal wiring toward maximizing ATP yield via the TCA–ETC axis, with ATP synthase as the central driver. Rather than relying on assumed objectives, ObjFind-M identifies the internal decision making that shapes cell physiology. Applied to CHO cells as a data-rich mammalian cell system, this inference of objectives through inverse modeling holds broad relevance, from optimizing biologics and cell and gene therapy production to deepening our knowledge and understanding of the drivers behind metabolic reprogramming in cancer and genetic disorders.

## Code and Data Availability

All the data used in this work is available in the Supplementary Data files. The ObjFind-M code is available at https://github.com/J-Morrissey/ObjFind-M.

## Author contributions

*Morrissey*: Conceptualization; Data curation; Formal analysis; Investigation; Methodology; Writing – original draft. *Monteiro*: Conceptualization; Writing – review and editing.

*Betenbaugh*: Funding acquisition; Supervision; Writing – review & editing. *Kontoravdi*: Funding acquisition; Supervision; Writing – review & editing.

## Acknowledgements

MM thanks the UK Biotechnology and Biological Sciences Research Council (BBSRC) and GSK for her studentship (BBSRC grant BB/I017011/1). This research was supported in part by NSF Award #2100800 to MJB. We are grateful to Max Mowbray for his valuable insights that contributed to this project.

## Competing interests

The authors declare no competing interests.

## Methods

### Methods table

| Data sources |  |  |
| --- | --- | --- |
| Fluxomics and metabolomics: CHO-DG44 | Templeton et al., 2013 | https://doi.org/10.1002/bit.24858 |
| Fluxomics and metabolomics: CHO-S with/without BCL2 overexpression | Templeton et al., 2014 | https://doi.org/10.1016/j.ymbe.2014.06.010 |
| Fluxomics and metabolomics: CHO-K1 with/without BCL2 overexpression | Templeton et al., 2017 | https://doi.org/10.1016/j.ymbe.2017.01.008 |
| Fluxomics and metabolomics: CHO-K1 in fed-batch and perfusion | Templeton et al., 2017 | https://doi.org/10.1016/j.ymbe.2017.09.010 |
| Fluxomics and metabolomics: CHO cells with/without ammonia supplementation | Pereira et al., 2018 | https://doi.org/10.1002/biot.201700518 |
| Fluxomics and metabolomics: CHO–Cum2 cells with/without induction of antibody expression | Sheikholeslami et al., 2013 | https://doi.org/10.1016/j.jbiotec.2013.01.025 |
| Fluxomics and metabolomics: CHO-K1 | Ahn & Antoniewicz, 2011 | https://doi.org/10.1016/j.ymbe.2011.07.002 |
| Fluxomics and metabolomics: CHO-K1 | Ahn & Antoniewicz, 2013 | https://doi.org/10.1016/j.ymbe.2012.10.001 |
| Fluxomics and metabolomics: CHO cells expressing zero, one or three anti-apoptotic gene(s) | Pereira, 2018 | https://ir.vanderbilt.edu/items/2aff3ac7-a0c1-4695-a40b-64ceb83d2378 |
| Fluxomics and metabolomics: CHO-K1 cells with attenuated GS promoter variants and selection conditions | Sacco et al., 2022 | https://doi.org/10.1002/bit.28084 |
| Fluxomics and metabolomics: CHO cells overexpressing PGC-1α | Sacco et al., 2023 | https://doi.org/10.1016/j.ymbe.2023.07.005 |
| Software and algorithms |  |  |
| ObjFind-M | This paper. | https://github.com/J-Morrissey/ObjFind-M |
| Python 3.0 | Python Software Foundation | https://www.python.org/downloads/ |
| COBRApy | Python Package | https://cobrapy.readthedocs.io/en/latest/ |
| Pyomo | Python Package | https://scipy.org/ |
| ipopt | COIN-OR | https://coin-or.github.io/Ipopt/ |

### ObjFind-M Data Inputs

#### Experimental inputs

The datasets used herein were selected from published literature of CHO cell culture that contained both ^13^C fluxomic metabolic flux analysis (MFA) and metabolite exchange data (see Methods Table). This consists of 42 experiments from 11 publications. We used reaction mapping tables to map the fluxomic MFA reactions from these datasets to the CHOmpact and iCHO2441 reactions, also performed in a previous method ^79^. Mapping tables for CHOmpact and iCHO2441 are provided in Supp. Data S1. One experiment was infeasible in iCHO2441 and four in CHOmpact and were removed from analysis of the respective model (Supp. Data S2). The raw input data is available in Supp. Data S7 and S8.

#### Stoichiometric networks

For this study, we used two stoichiometric networks: CHOmpact, a small-scale, model consisting of 144 reactions (205 accounting for splitting reversible reactions into two irreversible reactions) and 101 metabolites ^27^, which provides interpretability and a focus on core metabolism, and the CHO cell GEM iCHO2441 ^28^, a fusion of the iCHO2291 GEM ^35^ and the secretory pathway network from iCHO2048 ^31^. iCHO2441 contains 6337 reactions (8232 considering reversibility) and 4173 metabolites.

#### Mapping of ^13^C MFA Reactions to stoichiometric networks

Following collection of intracellular fluxes from ^13^C metabolic flux analysis (MFA) fluxomic data, the measured reactions must be mapped to their corresponding reactions in the CHO cell stoichiometric networks (CHOmpact and iCHO2441). The ^13^C MFA network is typically a lumped representation, where multiple elementary reactions are grouped into a single flux. As a result, one-to-one correspondence is rarely possible, and a systematic mapping framework is required.

The draft mapping is provided in Supplementary Data S1. To construct the mapping, the model reactions were treated as an electrical circuit, with “series” and “parallel” connections used to represent how elementary fluxes combine to form the net measured flux. Reactions in parallel were summed and treated as equivalent to a single reaction in series, after which the minimum flux across all series elements was taken to define the effective bound^80^. The following rules were applied to ensure consistency between ^13^C MFA fluxes and the mechanistic GEM:

- **Reaction directionality**: if a GEM reaction proceeded in the opposite orientation to the measured lumped flux, the flux value was multiplied by −1.
- **Stoichiometric scaling**: fluxes were multiplied by integer factors where necessary to match the stoichiometry of the lumped ^13^C reaction.
- **Compartmental redundancy**: where equivalent reactions occurred across multiple compartments, their contributions were summed to represent the single measured flux.

This mapping step established the correspondence between ^13^C MFA data and GEM reactions, allowing the measured fluxes to be implemented as constraints and objective terms in ObjFind-M.

### ObjFind-M Methods

#### ObjFind-M (“Inverse” Modeling)

We developed a bilevel optimization framework, ObjFind-M (Figure 1), as an adapted and expanded version of the Burgard and Maranas approach, originally applied to *E. Coli*, to infer the metabolic objective functions (or fitness function) of CHO cells using metabolomics and intracellular ^13^C flux data. Rather than assuming a fixed objective such as biomass maximization, ObjFind-M uses experimental metabolomics and ^13^C fluxomic to determine the set Coefficients of Fitness (*CoF*) that ensure model predictions match observed metabolic behavior.

At the outer level (or inference level) of the bilevel optimization, ObjFind-M treats the unknown fitness function as a set of coefficients that must be determined. The goal here is to find the combination of CoFs on candidate reactions that reproduces experimentally measured intracellular fluxes (from 13^C^ MFA fluxomics), as well as experimental growth rate and (where applicable) recombinant protein production. This level evaluates the agreement between predicted (*v*_*j*_) and measured 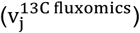 fluxes and adjusts the *CoF* in the inner level accordingly.

At the inner level (or FBA level), for each set of objective coefficients (*CoF*) proposed by the outer level, a standard FBA is performed. This level represents the CHO cell’s metabolism, calculating the flux distribution that would maximize the proposed objective function while satisfying steady-state mass balance, experimental metabolite uptake/secretion rates, and reaction capacity constraint. By solving the inner problem, we obtain the predicted metabolic state that the outer level then compares against the experimental data. The two levels are linked in the bilevel optimization as the outer level searches for the CoFs that make the inner level’s simulated behavior align close to the experimentally observed behavior. The ObjFind-M bilevel optimization formulation is written below:

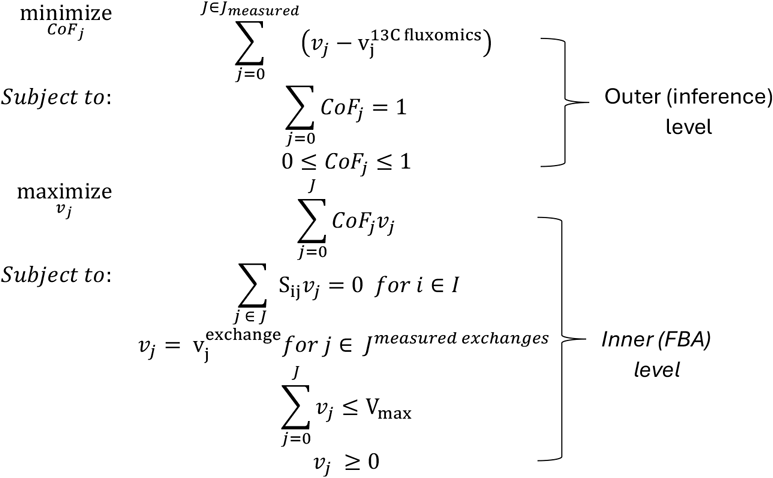

Where S_ij_ represents the stoichiometric coefficient of metabolite *i* in reaction 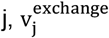 are the exchange rates of 20 amino acids, glucose and lactate. These are constrained to the median experimental values for those measured exchange reactions (Supp Data S7) in the datasets that correspond with the ^13^C MFA fluxomics (Supp Data S8) which are used as inputs to the outer level objective.

The outer level seeks to identify the (*CoF*_*j*_) that yields an FBA solution (*v*_*j*_) best matching the measured intracellular fluxes 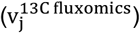. This is achieved by minimizing the squared deviation across all reactions for which intracellular flux measurements are available, using the median experimentally determined value from ^13^C fluxomics, mapped to the stoichiometric network using the mapping method outlined above. Experimentally determined growth rate and recombinant protein production (for producer cells) are also included within the outer level minimization, meaning ObjFind-M tries to solve metabolic behavior to achieve these values. Further information on the formulation of the outer level is found in Supp. Document A2.

Reversible reactions were split into two irreversible reactions to prevent negative flux and hence *v*_*j*_ must be greater than or equal to zero. The total flux sum was capped to mitigate the effects of thermodynamically infeasible cycles and therefore avoid unrealistic solutions.

This upper limit could be based on the minimal sum of flux ^81^, or a physiological capacity such as the enzymatic capacity constraint ^82^, however we selected a consistent upper limit (V_max_) across all experiments, with 10 and 100 mmol g^-1^ hr^-1^ chosen for CHOmpact and iCHO2441 respectively, due to relative size of each network.

#### Formulation of the Outer Level Objective

In the manuscript, the outer level objective is written as:

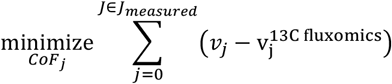

Once ^13^C MFA fluxes were mapped to GEM reactions, as outlined in S2 above, each mapped flux was assigned an **objective coefficient** to ensure balanced weighting in the optimization (not to be confused with CoF). In ObjFind-M, each ^13^C MFA reaction that is mapped to the GEM is assigned a total objective weight of 1. For a direct one-to-one (series) mapping, the single GEM reaction inherits the full weight. For cases where a ^13^C MFA reaction corresponds to several GEM reactions operating in parallel, the total weight is divided equally among the independent reactions. Specifically, each reaction is given an objective coefficient 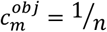, where *n* is the number of independent reactions in that mapping. For every ^13^C MFA reaction, the sum of objective coefficients across its mapped GEM reactions equals 1. In this way, ObjFind-M treats each measured ^13^C reaction equally in the optimization, without bias toward those that map to larger or more complex groups of reactions.

The outer level of ObjFind-M then formulates an optimization problem to minimize the deviation between predicted and experimental fluxes. The predicted fluxes are obtained from the inner FBA level given a trial set of coefficients of fitness (CoF). The objective function is:

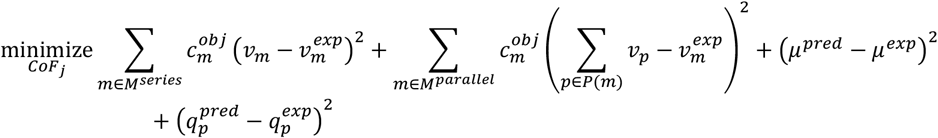

Here, *M*^*series*^ is the set of reactions with series mapping, *M*^*parallel*^ is the set of lumped reactions mapping to multiple GEM reactions, and *P*(*m*) denotes the reactions in parallel group *m*. The terms *μ* and *q*_*p*_ represent growth rate and recombinant protein productivity, respectively, which are also fitted in the outer objective. By minimizing this objective, ObjFind-M identifies the set of CoFs that best align the GEM-predicted fluxes with experimental data.

#### Inner Level Primal-Dual Reformulation

Solving ObjFind-M requires reformulating the inner level of the bilevel problem. The inner problem is a standard flux balance analysis (FBA) linear program, where a flux distribution is chosen to maximize a candidate objective function subject to stoichiometric and capacity constraints. Directly embedding this primal problem inside the outer level is computationally intractable. To address this, the inner FBA is reformulated into its dual representation^83^, which allows the bilevel system to be collapsed into a single-level optimization problem.

The primal problem is the standard FBA formulation:

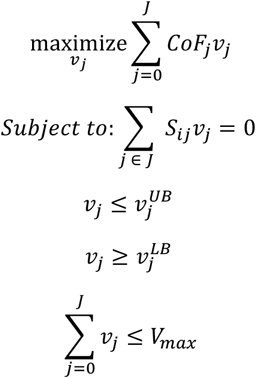

where *S*_*ij*_is the stoichiometric matrix, *v*_*j*_ are fluxes, and *CoF*_*j*_ are coefficients of the fitness function. The corresponding **dual problem** is:

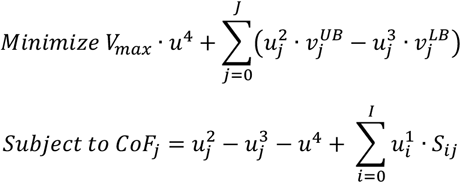

In the dual problem, the mass balance equations are associated with dual variables 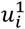, which are unrestricted in sign. Flux upper bounds introduce nonnegative duals 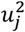, while flux lower bounds introduce nonnegative duals 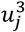. Finally, the global flux cap on the total sum of fluxes has a nonnegative dual variable *u*^4^.

The concept of strong duality guarantees that if the primal FBA has an optimal solution, so does the dual, and the two share the same optimal objective value. This property means that the primal and dual can only be simultaneously feasible at their optimal solutions. Therefore, by constructing a single optimization problem that incorporates both the primal feasibility constraints and the dual feasibility constraints, together with an equality constraint enforcing that the primal and dual objective values are identical, we collapse the bilevel system into one coherent formulation. In this way, any feasible solution to the combined problem is necessarily optimal for both the primal and dual, allowing ObjFind-M to solve for fitness coefficients while ensuring the inner FBA is optimally satisfied.

#### ObjFind-M Solution Methodology

The inferred fitness function from ObjFind-M is dependent on several factors, the first being the constraints applied to the metabolic model. In ObjFind-M, measured exchange rates of metabolites (20 common amino acids, glucose, and lactate) are constrained to their experimentally determined values within the inner level, meaning that these exchange rates will not appear within the fitness function. We instead focus on determining how the cell allocates the internal fluxes given these available resources ^4^. When comparing high and low producers, as well as when evaluating flux distributions, we also set growth rate and recombinant protein production in the inner/FBA level.

In the CHOmpact network, fluxes for NADP → NAD (F77) and NADH → NAD (F103) are set to zero to avoid “free” cofactor interconversions. Flux through F144, which defines the consumption of ATP toward active amino acid transport (ATP →), is also set to zero so that F104 remains the sole ATP demand reaction. All other reactions are left unconstrained apart from standard irreversibility settings, with biomass and recombinant protein production rates incorporated into the outer-level optimization.

Another factor impacting objective choice is the structure of the objective function itself. For the sake of simplicity, computational efficiency, as well as to find the importance of individual reactions, this paper focuses only on linear combinations of reactions to form the fitness function. To prevent any bias on any pre-determined objective, we allow all reactions to be assigned a CoF. The third factor affecting results is the choice of solver. Due to computational time, particularly for iCHO2441, we selected the local solver “ipopt” ^84^, which makes the problem solvable in a reasonable timeframe, but risks finding sub-optimal solutions. We observed that, for CHOmpact, after >100 restarts, only a handful of multiple optima were identified (Supp. Data S9). ObjFind-M was implemented and solved using Pyomo v6.6.1 ^85^.

#### Dual price

To complement the interpretation of the inferred metabolic objectives, we examined the *dual price* (or *shadow price*) of the stationary Lagrangian constraint associated with each reaction in the ObjFind-M formulation. In the KKT-based single-level reformulation of our bilevel optimization problem, this constraint mathematically defines coefficient *CoF*_*j*_ of a reaction in terms of the gradient of the Lagrangian with respect to the reaction flux *v*_*j*_. Conceptually, this ensures that *CoF*_*j*_ is consistent with the optimality conditions of the inner FBA problem.

The dual price on that constraint quantifies how sensitive the model’s solution is to enforcing this exact relationship between *CoF*_*j*_ and the flux *v*_*j*_,i.e., how essential the reaction *v*_*j*_ is to the structure of the optimal solution, because *CoF*_*j*_ is defined *through* that structure. While *CoF*_*j*_ gives the importance of each reaction to the network, the dual variable tells you which reactions are structurally critical for ensuring the objective fits the observed behavior. We computed the dual prices directly from the solution of the optimization problem using the ipopt solver. For each experiment, the dual price of the stationary constraint for reaction *j* was extracted from the solver’s Lagrange multipliers.

### General methods

#### Flux balance analysis (FBA)

FBA is an optimization problem to solve a constrained metabolic model. Constrained metabolic models are underdetermined since unknown reaction fluxes (*v*_*j*_) typically outnumber steady state mass balances and fluxes that can be determined using experimental data used to constrain the model. To solve for a flux distribution an objective function (*v*_*objective*_) is selected, through which flux is maximized (or minimized).

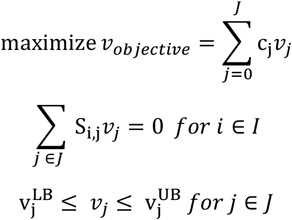

Where S_i,j_ is the stoichiometric coefficient of metabolite *i* in reaction *j, v*_*j*_ is the flux for reaction j and 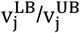 are the lower and upper bounds for reaction *j*. The metabolic objective can be constructed as a linear combination of reaction fluxes, with pre-determined weights c_j_ specifying each reaction’s contribution to the objective. In practice, this objective may be a single reaction (e.g., biomass, ATP, product secretion) or a composite that blends multiple cellular tasks (e.g., growth, energy, redox, product demand) ^86^. In our study, rather than pre-determining c_j_, we infer these coefficients from data.

#### Principal component analysis (PCA)

PCA was implemented to compare the inferred fitness function between high and low producers using sci-kit ^87^. A matrix was assembled with samples as rows and reactions CoF as columns, then each reaction column was z-scored (zero mean, unit variance) using scikit-learn’s StandardScaler. PCA was applied and the first two components were retained for visualization. Component loadings (pca.components) were extracted to rank reactions by their absolute contribution to PC1 and PC2.

#### Evaluation of commonly assumed objectives

To assess the predictive performance of widely used metabolic objectives for CHO cell models, we evaluated several candidate objectives within the iCHO2441 framework. FBA was carried out for the following objective functions and their corresponding reactions: biomass maximization (“biomass_cho”), recombinant protein maximization (“ICproduct_Final_demand”), ATP demand maximization (“DM_atp[c]”), Complex V maximization (“ATPS4m”), cytosolic NADPH minimization (all reactions producing metabolite “nadph[c]”), and minimization of the total flux sum (parsimonious FBA).

For these evaluations, the objective reactions being tested were left unconstrained, (e.g. growth rate was unconstrained during biomass maximization), while all exchange fluxes for amino acids, glucose, lactate, growth rate, and antibody production were fixed to the median experimentally measured value. During ObjFind-M, biomass, recombinant protein production, and ATP maintenance were left unconstrained, but during objective evaluation growth and recombinant protein production are constrained to the experimental value. ATP maintenance was constrained to a minimum value of 1.5 mmol g^-1^ hr^-1^, which is lower than reported for CHO ^80^, but maintained feasibility.

All optimizations were implemented in the COBRApy toolbox ^88^. Intracellular ^13^C fluxomics data were not used to constrain the model and were instead used as an independent dataset to evaluate the predictive accuracy of each objective, following the approach in ^28,79^. Performance was quantified using *root mean squared error (RMSE)* between the experimental ^13^C MFA flux value and model predicted flux.

#### Flux sampling

Flux sampling is an alternative method to predict flux distributions ^89^. Flux sampling, like FBA, uses steady state mass balances and constraints on reaction fluxes. However, it does not assume a metabolic objective, instead flux distributions are sampled from the solution space. Flux sampling was benchmarked as an alternative to FBA-based approaches. For this optGpSampler ^90^ was applied using the ‘model.sample’ function in the flux analysis submodule of COBRApy ^88^. The solution space was sampled 50,000,000 times, of which solutions were stored every 10,000 iterations, resulting in 5000 data-points per reaction. CycleFreeFlux ^91^ was then applied to the resulting flux distribution, and the median value was taken as a comparison to FBA-based approaches.

## Notes

### Competing Interest Statement

The authors have declared no competing interest.

## References

1. Feist, A.M., and Palsson, B.O. (2016). What do cells actually want? Genome Biol 16. 10.1186/s13059-016-0983-3.

2. Lin, D.-W., Khattar, S., and Chandrasekaran, S. (2025). Metabolic Objectives and Trade-Offs: Inference and Applications. Metabolites 15. 10.3390/metabo15020101.

3. Shoval, O., Sheftel, H., Shinar, G., Hart, Y., Ramote, O., Mayo, A., Dekel, E., Kavanagh, K., and Alon, U. (2012). Evolutionary trade-offs, pareto optimality, and the geometry of phenotype space. Science (1979) 336. 10.1126/science.1217405.

4. Islam, M.M., Shao, W., Batavia, M., Ford, R.M., Palsson, B.O., Nielsen, J., Maranas, C.D., Lee, S.Y., and Papin, J.A. (2025). Reframing the role of the objective function in its proper context for metabolic network modeling. Cell Syst 16, 101298. 10.1016/j.cels.2025.101298.

5. O’Brien, E.J., Monk, J.M., and Palsson, B.O. (2015). Using genome-scale models to predict biological capabilities. Preprint, 10.1016/j.cell.2015.05.019 https://doi.org/10.1016/j.cell.2015.05.019.

6. Feist, A.M., and Palsson, B.O. (2010). The biomass objective function. Preprint, 10.1016/j.mib.2010.03.003 https://doi.org/10.1016/j.mib.2010.03.003.

7. Bich, L., Pradeu, T., and Moreau, J.F. (2019). Understanding Multicellularity: The Functional Organization of the Intercellular Space. Front Physiol 10. 10.3389/fphys.2019.01170.

8. Magistretti, P.J., and Allaman, I. (2015). A Cellular Perspective on Brain Energy Metabolism and Functional Imaging. Preprint, 10.1016/j.neuron.2015.03.035 https://doi.org/10.1016/j.neuron.2015.03.035.

9. Hargreaves, M., and Spriet, L.L. (2020). Skeletal muscle energy metabolism during exercise. Preprint, 10.1038/s42255-020-0251-4 https://doi.org/10.1038/s42255-020-0251-4.

10. Orman, M.A., Mattick, J., Androulakis, I.P., Berthiaume, F., and Ierapetritou, M.G. (2012). Stoichiometry based steady-state hepatic flux analysis: Computational and experimental aspects. Preprint, 10.3390/metabo2010268 https://doi.org/10.3390/metabo2010268.

11. Bordbar, A., Monk, J.M., King, Z.A., and Palsson, B.O. (2014). Constraint-based models predict metabolic and associated cellular functions. Preprint, 10.1038/nrg3643 https://doi.org/10.1038/nrg3643.

12. Lewis, N.E., Nagarajan, H., and Palsson, B.O. (2012). Constraining the metabolic genotype-phenotype relationship using a phylogeny of in silico methods. Preprint, 10.1038/nrmicro2737 https://doi.org/10.1038/nrmicro2737.

13. Orth, J.D., Thiele, I., and Palsson, B.O. (2010). What is flux balance analysis? Preprint, 10.1038/nbt.1614 https://doi.org/10.1038/nbt.1614.

14. Schinn, S.M., Morrison, C., Wei, W., Zhang, L., and Lewis, N.E. (2021). Systematic evaluation of parameters for genome-scale metabolic models of cultured mammalian cells. Metab Eng 66. 10.1016/j.ymben.2021.03.013.

15. Gopalakrishnan, S., Johnson, W., Valderrama-Gomez, M.A., Icten, E., Tat, J., Ingram, M., Fung Shek, C., Chan, P.K., Schlegel, F., Rolandi, P., et al. (2024). COSMIC-dFBA: A novel multi-scale hybrid framework for bioprocess modeling. Metab Eng 82. 10.1016/j.ymben.2024.02.012.

16. Ramos, J.R.C., Oliveira, G.P., Dumas, P., and Oliveira, R. (2022). Genome-scale modeling of Chinese hamster ovary cells by hybrid semi-parametric flux balance analysis. Bioprocess Biosyst Eng 45. 10.1007/s00449-022-02795-9.

17. Lin, D.-W., Zhang, L., Zhang, J., and Chandrasekaran, S. (2025). Inferring metabolic objectives and trade-offs in single cells during embryogenesis. Cell Syst 16. 10.1016/j.cels.2024.12.005.

18. Park, S.-Y., Choi, D.-H., Song, J., Lakshmanan, M., Richelle, A., Yoon, S., Kontoravdi, C., Lewis, N.E., and Lee, D.-Y. (2024). Driving towards digital biomanufacturing by CHO genome-scale models. Trends Biotechnol 42, 1192–1203. 10.1016/j.tibtech.2024.03.001.

19. Ranpura, S., Maralingannavar, V., Gheorghe, A.-G., Ma, E., Morrissey, J., Betenbaugh, M.J., and Demirhan, D. (2025). Wheels turning: CHO cell modeling moves into a digital biomanufacturing era: Subtitle: CHO Metabolic Modeling. Comput Struct Biotechnol J 27, 2796–2813. 10.1016/j.csbj.2025.06.035.

20. Robinson, J.L., Kocabaş, P., Wang, H., Cholley, P.E., Cook, D., Nilsson, A., Anton, M., Ferreira, R., Domenzain, I., Billa, V., et al. (2020). An atlas of human metabolism. Sci Signal 13. 10.1126/scisignal.aaz1482.

21. Brunk, E., Sahoo, S., Zielinski, D.C., Altunkaya, A., Dräger, A., Mih, N., Gatto, F., Nilsson, A., Preciat Gonzalez, G.A., Aurich, M.K., et al. (2018). Recon3D enables a three-dimensional view of gene variation in human metabolism. Nat Biotechnol 36, 272–281. 10.1038/nbt.4072.

22. Burgard, A.P., and Maranas, C.D. (2003). Optimization-based framework for inferring and testing hypothesized metabolic objective functions. Biotechnol Bioeng 82. 10.1002/bit.10617.

23. Gianchandani, E.P., Oberhardt, M.A., Burgard, A.P., Maranas, C.D., and Papin, J.A. (2008). Predicting biological system objectives de novo from internal state measurements. BMC Bioinformatics 9. 10.1186/1471-2105-9-43.

24. Zhao, Q., Stettner, A.I., Reznik, E., Paschalidis, I.C., and Segrè, D. (2016). Mapping the landscape of metabolic goals of a cell. Genome Biol 17. 10.1186/s13059-016-0968-2.

25. Hart, Y., Sheftel, H., Hausser, J., Szekely, P., Ben-Moshe, N.B., Korem, Y., Tendler, A., Mayo, A.E., and Alon, U. (2015). Inferring biological tasks using Pareto analysis of high-dimensional data. Nat Methods 12. 10.1038/nmeth.3254.

26. Richelle, A., Kellman, B.P., Wenzel, A.T., Chiang, A.W.T., Reagan, T., Gutierrez, J.M., Joshi, C., Li, S., Liu, J.K., Masson, H., et al. (2021). Model-based assessment of amammalian cell metabolic functionalities using omics data. Cell Reports Methods 1. 10.1016/j.crmeth.2021.100040.

27. Jiménez del Val, I., Kyriakopoulos, S., Albrecht, S., Stockmann, H., Rudd, P.M., Polizzi, K.M., and Kontoravdi, C. (2023). CHOmpact: A reduced metabolic model of Chinese hamster ovary cells with enhanced interpretability. Biotechnol Bioeng 120. 10.1002/bit.28459.

28. Strain, B., Morrissey, J., Antonakoudis, A., and Kontoravdi, C. (2023). How reliable are Chinese hamster ovary (CHO) cell genome-scale metabolic models? Biotechnol Bioeng 120. 10.1002/bit.28366.

29. Park, J.U., Han, H.J., and Baik, J.Y. (2022). Energy metabolism in Chinese hamster ovary (CHO) cells: Productivity and beyond. Preprint, 10.1007/s11814-022-1062-y https://doi.org/10.1007/s11814-022-1062-y.

30. Coulet, M., Kepp, O., Kroemer, G., and Basmaciogullari, S. (2022). Metabolic Profiling of CHO Cells during the Production of Biotherapeutics. Preprint, 10.3390/cells11121929 https://doi.org/10.3390/cells11121929.

31. Gutierrez, J.M., Feizi, A., Li, S., Kallehauge, T.B., Hefzi, H., Grav, L.M., Ley, D., Baycin Hizal, D., Betenbaugh, M.J., Voldborg, B., et al. (2020). Genome-scale reconstructions of the mammalian secretory pathway predict metabolic costs and limitations of protein secretion. Nat Commun 11. 10.1038/s41467-01913867-y.

32. O’Brien, C.M., Mulukutla, B.C., Mashek, D.G., and Hu, W.S. (2020). Regulation of Metabolic Homeostasis in Cell Culture Bioprocesses. Preprint, 10.1016/j.tibtech.2020.02.005 https://doi.org/10.1016/j.tibtech.2020.02.005.

33. Martínez, V.S., Dietmair, S., Quek, L.E., Hodson, M.P., Gray, P., and Nielsen, L.K. (2013). Flux balance analysis of CHO cells before and after a metabolic switch from lactate production to consumption. Biotechnol Bioeng 110. 10.1002/bit.24728.

34. Vazquez, A., and Oltvai, Z.N. (2016). Macromolecular crowding explains overflow metabolism in cells. Sci Rep 6. 10.1038/srep31007.

35. Yeo, H.C., Hong, J., Lakshmanan, M., and Lee, D.Y. (2020). Enzyme capacity-based genome scale modelling of CHO cells. Metab Eng 60, 138–147. 10.1016/j.ymben.2020.04.005.

36. Morrissey, J., Cankorur-Cetinkaya, A., Grassi, L., Harwood-Stamper, A.J., Welsh, J., and Kontoravdi, C. (2025). Nicotinamide reverses the Warburg effect in Chinese hamster ovary cell culture. bioRxiv. 10.1101/2025.06.18.660349.

37. Luengo, A., Li, Z., Gui, D.Y., Sullivan, L.B., Zagorulya, M., Do, B.T., Ferreira, R., Naamati, A., Ali, A., Lewis, C.A., et al. (2021). Increased demand for NAD+ relative to ATP drives aerobic glycolysis. Mol Cell 81. 10.1016/j.molcel.2020.12.012.

38. Locasale, J.W., and Cantley, L.C. (2011). Metabolic flux and the regulation of mammalian cell growth. Preprint, 10.1016/j.cmet.2011.07.014 https://doi.org/10.1016/j.cmet.2011.07.014.

39. Pereira, S., Kildegaard, H.F., and Andersen, M.R. (2018). Impact of CHO Metabolism on Cell Growth and Protein Production: An Overview of Toxic and Inhibiting Metabolites and Nutrients. Preprint, 10.1002/biot.201700499 https://doi.org/10.1002/biot.201700499.

40. Kirsch, B.J., Bennun, S. V., Mendez, A., Johnson, A.S., Wang, H., Qiu, H., Li, N., Lawrence, S.M., Bak, H., and Betenbaugh, M.J. (2022). Metabolic analysis of the asparagine and glutamine dynamics in an industrial Chinese hamster ovary fed-batch process. Biotechnol Bioeng 119. 10.1002/bit.27993.

41. Torres, M., Elvin, M., Betts, Z., Place, S., Gaffney, C., and Dickson, A.J. (2021). Metabolic profiling of Chinese hamster ovary cell cultures at different working volumes and agitation speeds using spin tube reactors. Biotechnol Prog 37. 10.1002/btpr.3099.

42. Carinhas, N., Duarte, T.M., Barreiro, L.C., Carrondo, M.J.T., Alves, P.M., and Teixeira, A.P. (2013). Metabolic signatures of GS-CHO cell clones associated with butyrate treatment and culture phase transition. Biotechnol Bioeng 110, 3244–3257. 10.1002/bit.24983.

43. Mráček, T., Drahota, Z., and Houštěk, J. (2013). The function and the role of the mitochondrial glycerol-3-phosphate dehydrogenase in mammalian tissues. Preprint, 10.1016/j.bbabio.2012.11.014 https://doi.org/10.1016/j.bbabio.2012.11.014.

44. Mulukutla, B.C., Mitchell, J., Geoffroy, P., Harrington, C., Krishnan, M., Kalomeris, T., Morris, C., Zhang, L., Pegman, P., and Hiller, G.W. (2019). Metabolic engineering of Chinese hamster ovary cells towards reduced biosynthesis and accumulation of novel growth inhibitors in fed-batch cultures. Metab Eng 54. 10.1016/j.ymben.2019.03.001.

45. Vatrinet, R., Leone, G., De Luise, M., Girolimetti, G., Vidone, M., Gasparre, G., and Porcelli, A.M. (2017). The α-ketoglutarate dehydrogenase complex in cancer metabolic plasticity. Cancer Metab 5. 10.1186/s40170-017-0165-0.

46. Reed, L.J., and Hackert, M.L. (1990). Structure-function relationships in dihydrolipoamide acyltransferases. Preprint, 10.1016/s0021-9258(19)38795-2 https://doi.org/10.1016/s0021-9258(19)38795-2.

47. Sellick, C.A., Croxford, A.S., Maqsood, A.R., Stephens, G.M., Westerhoff, H. V., Goodacre, R., and Dickson, A.J. (2015). Metabolite profiling of CHO cells: Molecular reflections of bioprocessing effectiveness. Biotechnol J 10. 10.1002/biot.201400664.

48. Selvarasu, S., Ho, Y.S., Chong, W.P.K., Wong, N.S.C., Yusufi, F.N.K., Lee, Y.Y., Yap, M.G.S., and Lee, D.Y. (2012). Combined in silico modeling and metabolomics analysis to characterize fed-batch CHO cell culture. Biotechnol Bioeng 109. 10.1002/bit.24445.

49. Verhagen, N., Teleki, A., Heinrich, C., Schilling, M., Unsöld, A., and Takors, R. (2020). S-adenosylmethionine and methylthioadenosine boost cellular productivities of antibody forming Chinese hamster ovary cells. Biotechnol Bioeng 117. 10.1002/bit.27484.

50. Templeton, N., Lewis, A., Dorai, H., Qian, E.A., Campbell, M.P., Smith, K.D., Lang, S.E., Betenbaugh, M.J., and Young, J.D. (2014). The impact of anti-apoptotic gene Bcl-2Δ expression on CHO central metabolism. Metab Eng 25. 10.1016/j.ymben.2014.06.010.

51. Sengupta, N., Rose, S.T., and Morgan, J.A. (2011). Metabolic flux analysis of CHO cell metabolism in the late non-growth phase. Biotechnol Bioeng 108. 10.1002/bit.22890.

52. Ahn, W.S., and Antoniewicz, M.R. (2011). Metabolic flux analysis of CHO cells at growth and non-growth phases using isotopic tracers and mass spectrometry. Metab Eng 13. 10.1016/j.ymben.2011.07.002.

53. Woo Suk, A., and Antoniewicz, M.R. (2013). Parallel labeling experiments with [1,2-13C]glucose and [U-13C]glutamine provide new insights into CHO cell metabolism. Metab Eng 15. 10.1016/j.ymben.2012.10.001.

54. Herzig, S., Raemy, E., Montessuit, S., Veuthey, J.L., Zamboni, N., Westermann, B., Kunji, E.R.S., and Martinou, J.C. (2012). Identification and functional expression of the mitochondrial pyruvate carrier. Science (1979) 336. 10.1126/science.1218530.

55. Brooks, G.A. (2018). The Science and Translation of Lactate Shuttle Theory. Preprint, 10.1016/j.cmet.2018.03.008 https://doi.org/10.1016/j.cmet.2018.03.008.

56. Luginsland, M., Kontoravdi, C., Racher, A., Jaques, C., and Kiparissides, A. (2024). Elucidating lactate metabolism in industrial CHO cultures through the combined use of flux balance and principal component analyses. Biochem Eng J 202. 10.1016/j.bej.2023.109184.

57. Yusufi, F.N.K., Lakshmanan, M., Ho, Y.S., Loo, B.L.W., Ariyaratne, P., Yang, Y., Ng, S.K., Tan, T.R.M., Yeo, H.C., Lim, H.L., et al. (2017). Mammalian Systems Biotechnology Reveals Global Cellular Adaptations in a Recombinant CHO Cell Line. Cell Syst 4. 10.1016/j.cels.2017.04.009.

58. Wilkens, C.A., and Gerdtzen, Z.P. (2015). Comparative metabolic analysis of CHO cell clones obtained through cell engineering, for IgG productivity, growth and cell longevity. PLoS One 10. 10.1371/journal.pone.0119053.

59. Templeton, N., Xu, S., Roush, D.J., and Chen, H. (2017). 13C metabolic flux analysis identifies limitations to increasing specific productivity in fed-batch and perfusion. Metab Eng 44. 10.1016/j.ymben.2017.09.010.

60. Huang, Z., Xu, J., Yongky, A., Morris, C.S., Polanco, A.L., Reily, M., Borys, M.C., Li, Z.J., and Yoon, S. (2020). CHO cell productivity improvement by genome-scale modeling and pathway analysis: Application to feed supplements. Biochem Eng J 160. 10.1016/j.bej.2020.107638.

61. Chong, W.P.K., Reddy, S.G., Yusufi, F.N.K., Lee, D.Y., Wong, N.S.C., Heng, C.K., Yap, M.G.S., and Ho, Y.S. (2010). Metabolomics-driven approach for the improvement of Chinese hamster ovary cell growth: Overexpression of malate dehydrogenase II. J Biotechnol 147. 10.1016/j.jbiotec.2010.03.018.

62. Templeton, N., Dean, J., Reddy, P., and Young, J.D. (2013). Peak antibody production is associated with increased oxidative metabolism in an industrially relevant fed-batch CHO cell culture. Biotechnol Bioeng 110. 10.1002/bit.24858.

63. Dean, J., and Reddy, P. (2013). Metabolic analysis of antibody producing CHO cells in fed-batch production. Biotechnol Bioeng 110. 10.1002/bit.24826.

64. Zhao, F., Wan, Y., Nie, L., Jiao, J., Gao, D., Sun, Y., Chen, Z., Shi, Y., Yang, J., Pan, J., et al. (2023). 1H NMR-based process understanding and biochemical marker identification methodology for monitoring CHO cell culture process during commercial-scale manufacturing. Biotechnol J 18. 10.1002/biot.202200616.

65. Eriksson, A., Richelle, A., Trygg, J., Scholze, S., Pijeaud, S., Antti, H., Zehe, C., Surowiec, I., and Jonsson, P. (2025). Time-Resolved Hierarchical Modeling Highlights Metabolites Influencing Productivity and Cell Death in Chinese Hamster Ovary Cells. Biotechnol J 20, e202400624. 10.1002/biot.202400624.

66. Calmels, C., Arnoult, S., Ben Yahia, B., Malphettes, L., and Andersen, M.R. (2019). Application of a genome-scale model in tandem with enzyme assays for identification of metabolic signatures of high and low CHO cell producers. Metab Eng Commun 9. 10.1016/j.mec.2019.e00097.

67. Barberi, G., Benedetti, A., Diaz-Fernandez, P., Sévin, D.C., Vappiani, J., Finka, G., Bezzo, F., and Facco, P. (2025). Productive CHO cell lines selection in biopharm process development through machine learning on metabolomic dynamics. AIChE Journal 71, e18602. 10.1002/aic.18602.

68. Lopaschuk, G.D., Ussher, J.R., Folmes, C.D.L., Jaswal, J.S., and Stanley, W.C. (2010). Myocardial fatty acid metabolism in health and disease. Preprint, 10.1152/physrev.00015.2009 https://doi.org/10.1152/physrev.00015.2009.

69. Schaub, J., Clemens, C., Schorn, P., Hildebrandt, T., Rust, W., Mennerich, D., Kaufmann, H., and Schulz, T.W. (2010). CHO gene expression profiling in biopharmaceutical process analysis and design. Biotechnol Bioeng 105. 10.1002/bit.22549.

70. Priem, B., Cai, X., Hong, Y.-J., Gilmore, K., Deng, Z., Chen, S., Naik, H.M., Betenbaugh, M.J., and Antoniewicz, M.R. (2025). Modulating fatty acid metabolism and composition of CHO cells by feeding high levels of fatty acids complexed using methyl-beta-cyclodextrin. Metab Eng 91, 158–169. 10.1016/j.ymben.2025.04.005.

71. Chong, W.P.K., Thng, S.H., Hiu, A.P., Lee, D.Y., Chan, E.C.Y., and Ho, Y.S. (2012). LC-MS-based metabolic characterization of high monoclonal antibody-producing Chinese hamster ovary cells. Biotechnol Bioeng 109. 10.1002/bit.24580.

72. García Sánchez, C.E., and STorres áez, R.G. (2014). Comparison and analysis of objective functions in flux balance analysis. Biotechnol Prog 30. 10.1002/btpr.1949.

73. Templeton, N., Smith, K.D., McAtee-Pereira, A.G., Dorai, H., Betenbaugh, M.J., Lang, S.E., and Young, J.D. (2017). Application of 13C flux analysis to identify high-productivity CHO metabolic phenotypes. Metab Eng 43. 10.1016/j.ymben.2017.01.008.

74. McAtee Pereira, A.G., Walther, J.L., Hollenbach, M., and Young, J.D. (2018). 13C Flux Analysis Reveals that Rebalancing Medium Amino Acid Composition can Reduce Ammonia Production while Preserving Central Carbon Metabolism of CHO Cell Cultures. Biotechnol J 13. 10.1002/biot.201700518.

75. Sheikholeslami, Z., Jolicoeur, M., and Henry, O. (2013). Probing the metabolism of an inducible mammalian expression system using extracellular isotopomer analysis. J Biotechnol 164. 10.1016/j.jbiotec.2013.01.025.

76. McAtee Pereira, A.G. (2018). 13C metabolic flux analysis of industrial Chinese hamster ovary (CHO) cell cultures.

77. Sacco, S.A., Tuckowski, A.M., Trenary, I., Kraft, L., Betenbaugh, M.J., Young, J.D., and Smith, K.D. (2022). Attenuation of glutamine synthetase selection marker improves product titer and reduces glutamine overflow in Chinese hamster ovary cells. Biotechnol Bioeng 119. 10.1002/bit.28084.

78. Sacco, S.A., McAtee Pereira, A.G., Trenary, I., Smith, K.D., Betenbaugh, M.J., and Young, J.D. (2023). Overexpression of peroxisome proliferator-activated receptor γ co-activator-1α (PGC-1α) in Chinese hamster ovary cells increases oxidative metabolism and IgG productivity. Metab Eng 79. 10.1016/j.ymben.2023.07.005.

79. Morrissey, J., Barberi, G., Strain, B., Facco, P., and Kontoravdi, C. (2025). NEXT-FBA: A hybrid stoichiometric/data-driven approach to improve intracellular flux predictions. Metab Eng. 10.1016/j.ymben.2025.03.010.

80. Széliová, D., Štor, J., Thiel, I., Weinguny, M., Hanscho, M., Lhota, G., Borth, N., Zanghellini, J., Ruckerbauer, D.E., and Rocha, I. (2021). Inclusion of maintenance energy improves the intracellular flux predictions of CHO. PLoS Comput Biol 17. 10.1371/journal.pcbi.1009022.

81. Lewis, N.E., Hixson, K.K., Conrad, T.M., Lerman, J.A., Charusanti, P., Polpitiya, A.D., Adkins, J.N., Schramm, G., Purvine, S.O., Lopez-Ferrer, D., et al. (2010). Omic data from evolved E. coli are consistent with computed optimal growth from genome-scale models. Mol Syst Biol 6. 10.1038/msb.2010.47.

82. Adadi, R., Volkmer, B., Milo, R., Heinemann, M., and Shlomi, T. (2012). Prediction of microbial growth rate versus biomass yield by a metabolic network with kinetic parameters. PLoS Comput Biol 8, e1002575. 10.1371/journal.pcbi.1002575.

83. Bertsimas, D., and Tsitsiklis, J. (1997). Introduction to Linear Optimization (Athena Scientific Series in Optimization and Neural Computation, 6). Book.

84. Wächter, A., and Biegler, L.T. (2006). Line search filter methods for nonlinear programming: Motivation and global convergence. SIAM Journal on Optimization 16. 10.1137/S1052623403426556.

85. Hart, W.E., Watson, J.P., and Woodruff, D.L. (2011). Pyomo: Modeling and solving mathematical programs in Python. Math Program Comput 3. 10.1007/s12532-011-0026-8.

86. Morrissey, J., Strain, B., and Kontoravdi, C. (2024). Flux Balance Analysis of Mammalian Cell Systems. In Methods in Molecular Biology 10.1007/978-1-0716-3718-0_9.

87. Pedregosa, F., Varoquaux, G., Gramfort, A., Michel, V., Thirion, B., Grisel, O., Blondel, M., Prettenhofer, P., Weiss, R., Dubourg, V., et al. (2011). Scikit-learn: Machine learning in Python. Journal of Machine Learning Research 12.

88. Ebrahim, A., Lerman, J.A., Palsson, B.O., and Hyduke, D.R. (2013). COBRApy: COnstraints-Based Reconstruction and Analysis for Python. BMC Syst Biol 7. 10.1186/1752-0509-7-74.

89. Herrmann, H.A., Dyson, B.C., Vass, L., Johnson, G.N., and Schwartz, J.M. (2019). Flux sampling is a powerful tool to study metabolism under changing environmental conditions. NPJ Syst Biol Appl 5. 10.1038/s41540-019-0109-0.

90. Megchelenbrink, W., Huynen, M., and Marchiori, E. (2014). optGpSampler: An improved tool for uniformly sampling the solution-space of genome-scale metabolic networks. PLoS One 9. 10.1371/journal.pone.0086587.

91. Desouki, A.A., Jarre, F., Gelius-Dietrich, G., and Lercher, M.J. (2015). CycleFreeFlux: Efficient removal of thermodynamically infeasible loops from flux distributions. Bioinformatics 31, 2159–2165. 10.1093/bioinformatics/btv096.

